# A computational model of the two dentate gyrus blades

**DOI:** 10.64898/2026.09.01.748300

**Authors:** Viktor Studenyak, Jürgen Jost, Christian F. Doeller, Andrej Bicanski

## Abstract

The Dentate Gyrus (DG) is a key part of the hippocampus, and damage to the DG produces a wide range of pathologies, including overgeneralization of contexts, affective dysregulation (Anacker et al., 2018), and epileptogenic effects (Sloviter, 1994). The canonical model of the DG focuses on pattern separation for subsequent memory storage in the hippocampal subfield CA3. Experimental results challenge the singular focus on pattern separation and extend the function of the DG to the precise binding of objects and events to space, and the integration of information across episodes. Recent studies suggest that pattern separation and integration preferentially rely on distinct DG blades, with the suprapyramidal and infrapyramidal blades biased toward separation and integration, respectively. Here, we propose the first computational model that accounts for this distinction: an exemplar-based k-WTA architecture in the suprapyramidal DG (DG_SUP_) supports pattern separation and episode-specific representations, whereas an architecture with gradual heterosynaptic plasticity in the infrapyramidal DG (DG_INF_) supports integration of patterns across episodes. Both coding regimes are tested with two datasets: MNIST and neurally plausible entorhinal cortex inputs, thus suggesting some domain generality. Using the entorhinal cortex inputs, the two blades form place fields that either remap or maintain a stable code, consistent with experimental results. Novel inputs, including novel digit classes and novel spatial episodes, are incorporated through a neurogenesis-inspired turnover and recruitment mechanism. The two processing streams allow for a comparison of ongoing experience with the generalized expectations formed through integration across episodes. This yields prediction errors that can drive the storage of poorly predicted memories and the forgetting of well-predicted memories. The differential processing across the DG could thus aid in the iterative construction of spatial cognitive maps that encode location-dependent expectations, while at the same time preserving individual episodic memory traces. These functions are accomplished with biologically plausible learning regimes and widen the scope of DG computation beyond its well-established role in pattern separation.

## Introduction

The hippocampal formation is crucial for memory formation, memory recall, and spatial navigation (Eichenbaum, 2000; Tulving, 2002). Marr’s seminal theory (Marr, 1971) suggested distinct roles in memory processes for the subregions of the hippocampus. In this account, the dentate gyrus (DG) plays a key role in pattern separation through an expansion in the number of cells and increased sparsity of the population code. This reduces the similarity of input patterns and prevents interference between overlapping memories in CA3. Subsequent modeling revealed that DG-mediated pattern separation can greatly facilitate memory encoding in CA3 (Treves and Rolls, 1994), and this view of the DG still constitutes the state of the art, and has been modeled repeatedly (Gibson et al., 1991; Kesner and Rolls, 2015; McClelland et al., 1995; Myers and Scharfman, 2011, 2009; O’Reilly and McClelland, 1994; Santhakumar et al., 2005).

Multiple experimental studies confirm that the dentate gyrus supports pattern separation at both behavioral and neural levels (Gilbert et al., 2001; Leutgeb et al., 2007; Neunuebel and Knierim, 2014). Thanks to improved experimental and machine-learning methods for distinguishing DG cell types, it has been shown that granule cells stay sparse and stable, with single place fields, whereas mossy cells remap globally and express multiple fields (Senzai and Buzsáki, 2017). An in vivo calcium imaging study further demonstrated that DG granule cells stay stable for days and only partially remap between familiar and novel environments, whereas CA3 place fields remap across days and in different environments (Hainmueller and Bartos, 2018). Together, this evidence suggests important departures from the standard account of pure pattern separation in the DG and suggests a potential role in episodic integration of memories to build a stable code of the environment (Hainmueller and Bartos, 2020).

The dentate gyrus is also one of only two locations in the adult brain with confirmed adult neurogenesis (Altman and Das, 1965; Eriksson et al., 1998). Neurogenesis appears to facilitate behavioral pattern separation (Clelland et al., 2009) and adult-born granule cells (abGCs) are involved in the discrimination of similar memories, whereas mature granule cells remain essential for pattern completion (Nakashiba et al., 2012). Moreover, abGCs seem to directly modulate mature granule cells (mGCs) depending on the origin of the inputs. In the suprapyramidal blade, activity of abGCs receiving lateral perforant-path input suppresses mGC activity, whereas in the infrapyramidal blade, abGCs that are predominantly driven by the medial path excite mGCs through extrasynaptic glutamatergic signalling (Luna et al., 2019). Facilitation of neurogenesis through genetic stem-cell expansion confirms the difference in the modulation of mGCs by abGCs, showing that enhanced neurogenesis improves memory discrimination and precision (Berdugo-Vega et al., 2021). Consistent with this, LEC preferentially projects to the suprapyramidal blade, whereas MEC preferentially targets the infrapyramidal blade (Canto et al., 2008; Witter et al., 2017). These anatomical and functional asymmetries have led to the hypothesis of two parallel processing streams: a pattern separation stream (spanning the lateral entorhinal cortex, the suprapyramidal DG, distal CA3 and proximal CA1) and an integration stream (spanning the medial entorhinal cortex, the infrapyramidal DG, proximal CA3 and distal CA1) (Berdugo-Vega et al., 2023).

The two-stream proposal has hitherto not been explored computationally. Most computational DG models concentrate on neurogenesis as the main distinctive feature of the DG and/or pattern separation. Existing models have explored additive and turnover neurogenesis, showing benefits for continual learning, reduction of memory interference and improved memory capacity (Chambers et al., 2004; Crick and Miranker, 2006; Wiskott et al., 2006; Aimone et al., 2009; Appleby and Wiskott, 2009; Weisz and Argibay, 2009; Appleby et al., 2011), as well as mechanisms such as the GABA-switch in the abGCs (Ge et al., 2006) that enable the encoding of novel patterns (Gozel and Gerstner, 2021).

While these influential models have significantly increased our understanding of DG function, to date, computational approaches have not addressed blade-specific differences, leaving the precise roles of the two DG blades elusive. Here, we propose the first computational model that investigates this distinction, in line with recent experimental work (Berdugo-Vega et al., 2021; Luna et al., 2019) and the proposed two-stream hypothesis (Berdugo-Vega et al., 2023). In our model, the suprapyramidal blade participates in storing distinct episodic memories through pattern separation and one-shot learning. Conversely, the infrapyramidal blade integrates information across episodes (learning at a slower rate) to form generalized expectations across episodes, eventually forming a cognitive-map-like representation of a familiar environment. These integrated representations form a cognitive-map-like representation that generates predictions about ongoing experience in an allocentric spatial reference frame, here implemented as allocentric object-vector relations anchored to the animal’s position with object identities and random spatial neurons. We propose that this dual slow/fast coding model of the DG can facilitate the comparison between new and old episodes with these learned expectations, enabling the computation of prediction errors that drive both the storage of poorly predicted (novel) memories and the forgetting of well-predicted (redundant) ones. This would allow the hippocampal system to free up neuronal resources and enable the model to iteratively build a spatial cognitive map for a familiar environment, one episode at a time.

We show that, based on different learning rules, the suprapyramidal and infrapyramidal blades of the DG perform complementary computations, separating inputs in the suprapyramidal blade and integrating them in the infrapyramidal blade. This integration happens across episodes, while pattern separation across locations remains preserved. We further show the ability of the network to encode novel inputs, including novel MNIST classes and novel spatial episodes, using neurogenesis-inspired recruitment and, in DG_INF_, a GABA-switch-like maturation mechanism.

## Results

### Model architecture and inputs

To explore the hypothesis that the two dentate gyrus (DG) blades support complementary encoding regimes, we implement related but functionally distinct architectures with different plasticity mechanisms. The suprapyramidal blade network consists of two layers: the entorhinal cortex (EC) input layer and the suprapyramidal dentate gyrus (DG_SUP_) layer (Fig. 1a). The infrapyramidal blade network additionally has an interneuron layer, forming a three-layer network (EC–DG_INF_–interneurons; Fig. 1b). A full model description is provided in the Methods.

**Figure 1.**
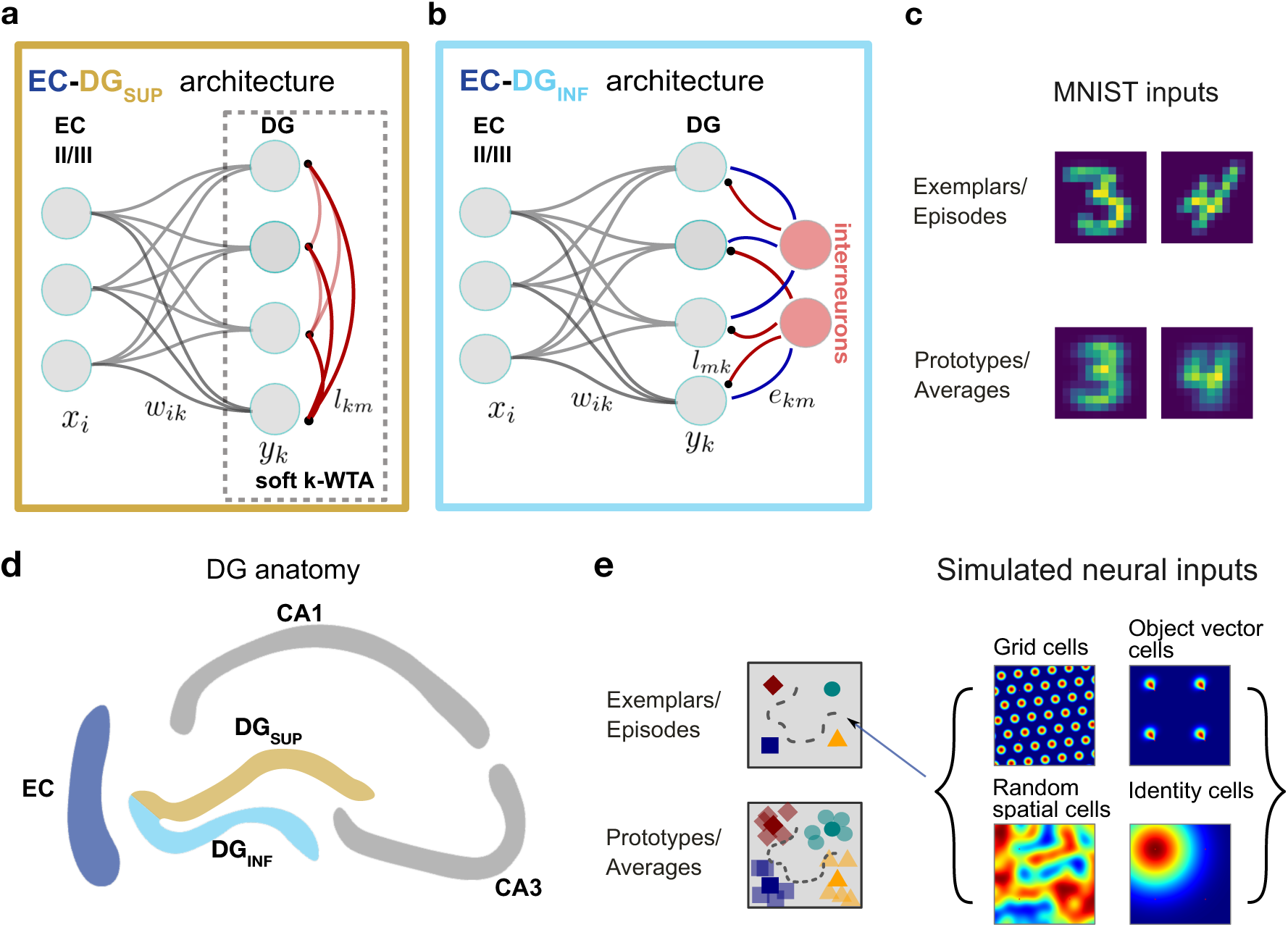
Dentate gyrus model architecture and input datasets. **a-b**: Model architectures for the two dentate gyrus (DG) blades. **a:** The suprapyramidal blade model, DG_SUP_, is implemented as an EC–DG network with learned feedforward weights and competitive inhibition, supporting rapid exemplar-like encoding. **b:** The infrapyramidal blade model, DG_INF_, includes an additional interneuron layer, forming an EC–DG–interneuron network that supports slower integration across related inputs. Yellow and blue indicate the DG_SUP_ and DG_INF_ processing streams, respectively. **c:** MNIST input dataset. Individual handwritten digit instances are used as exemplar-like episode analogues. The class averages as class prototypes are shown here, but are not part of the inputs. **d:** Schematic of the hippocampal formation, highlighting the DG blades and downstream hippocampal regions considered in the model. **e:** Simulated EC input dataset. Episodes are generated as trajectories through a 2D environment containing objects. Across episodes, object positions are slightly perturbed, allowing mean object locations to define integrated representations. Averaged object locations are shown, but are not part of the inputs. Example rate maps are shown for the simulated EC populations: grid cells, object-vector cells, random spatial cells, and identity cells.

The suprapyramidal blade is modeled as a feedforward EC–DG network with DG neurons that have no explicit intrinsic dynamics. In the model, learning is implemented via an exemplar-based competitive recruitment mechanism with k-Winner-Take-All (k-WTA) selection and soft activity readout, where neuron activity scales continuously with similarity to the stored exemplar rather than being binary. This makes it unnecessary to implement a separate interneuron layer as in DG_INF_. The k-WTA mechanism captures competitive allocation among stored exemplars. However, in DG_INF_ the presence of the IN layer allows for the modeling of the GABA-switch mechanism (see below). For each input, the similarity to existing representations is computed. If the maximal similarity is lower than the selectivity threshold *_θSUP_*, a previously uncommitted neuron is recruited. EC→DG synaptic weights are then imprinted in a one-shot manner, storing the input as an exemplar. Winner neurons whose similarity is at least *θ*_SUP_ undergo a fractional weight update with step size *η*_reuse_, allowing limited integration of highly similar inputs. Competition is implemented through learned lateral inhibition and k-WTA winner selection, whereas DG activity used for readout is computed as a graded soft similarity response over committed neurons. This architecture supports rapid exemplar storage and strong pattern separation of the inputs.

The infrapyramidal blade network (DG_INF_) consists of an EC input layer, a DG excitatory population, and an explicit inhibitory interneuron layer mediating lateral inhibition. Synaptic weights from EC to DG are plastic, whereas connections between DG neurons and interneurons are fixed. EC→DG synapses are trained using a heterosynaptic plasticity rule (Chistiakova et al., 2014; Zenke and Gerstner, 2017; Gozel and Gerstner, 2021), in which potentiation and depression are balanced by additional heterosynaptic terms, which enforce competition and homeostasis. In contrast to DG_SUP_, learning in DG_INF_ is slow and requires repeated exposure to the input patterns. This is implemented via a lower learning rate and multiple training epochs. This leads to gradual integration of similar inputs. We primarily view these as reflecting similar experiences, although they may also correspond to repeated reactivation of patterns during hippocampal replay (Wilson and McNaughton, 1994; Skaggs and McNaughton, 1996; Ólafsdóttir et al., 2018).

Functionally, the DG_SUP_ architecture implements pattern separation, potentially supported by inhibitory interactions between adult-born and mature granule cells in the suprapyramidal blade. Conversely, the DG_INF_ architecture promotes integration across patterns, potentially supported by excitatory interactions in this subnetwork (Luna et al., 2019; Berdugo-Vega et al., 2021, 2023). We evaluate the model using two input datasets: the MNIST handwritten digits and simulated entorhinal cortex (EC) population activity (Fig. 1c and 1e). MNIST enables direct visualization of receptive fields and provides a more readily interpretable analogy to episodic inputs. The simulated EC dataset is more biologically grounded and enables comparison to experimental observations. Moreover, consistent behavior across both datasets suggests that the proposed coding principles generalize across spatial and non-spatial domains.

For the simulated EC dataset, we generate population activity as an agent traverses an environment containing objects (Fig. 1e, left). We simulate four cell populations grouped into two anatomical systems. The medial entorhinal cortex (MEC) includes grid cells (GCs) (Hafting et al., 2005) and object-vector cells (OVCs) (Bicanski and Burgess, 2018; Høydal et al., 2019) (Fig. 1e, right). The lateral entorhinal cortex (LEC) includes random spatial (RS) cells and identity cells (IDCs) (Deshmukh and Knierim, 2011; Tsao et al., 2013) (Fig. 1e, right). We note that the biological two-stream hypothesis assumes a gradient of LEC and MEC routing to different DG blades (with some overlap). However, to isolate the contribution of blade-specific learning dynamics, we provide both subnetworks with the same combined EC input population. Thus, the functional dissociation between DG_SUP_ and DG_INF_ arises primarily from their different learning rules and circuit architectures, rather than from differences in the input statistics provided to each blade. An episode is defined as the sequence of EC population activity along one traversal through a fixed environment, with a particular object configuration and random spatial cell realization (Hasselmo, 2009, 2012; Bicanski, 2025). Similar episodes are generated within the same environment by slightly perturbing object positions and resampling random spatial cell tunings, which we assume provide an episode-specific contextual signal that may reflect changes in internal or motivational state (Kennedy and Shapiro, 2009). Repeated object displacements within the same environment allow estimation of mean object positions, corresponding to prototypical or expected locations. By contrast, distinct environments are generated by altering geometry and/or object configurations, accompanied by broader EC remapping including grid-cell rotations and shifts (Fyhn et al., 2007).

To build intuition, we can establish the following analogy between MNIST and EC inputs. Individual episodes can be analogized to specific digit instances or object configurations, whereas integrated representations might correspond to digit prototypes or expected object locations (Figure S1a).

### Pattern separation in DG_SUP_ and pattern integration in DG_INF_

We first trained both networks on MNIST inputs. For each input, we computed the activity-weighted average of receptive fields of active neurons and reshaped it into a 12 × 12 image (Fig. 2a, left). We observe that the DG_SUP_ representations closely match individual handwritten digits (Fig. 2a, middle), whereas DG_INF_ representations resemble class-averaged prototypes (Fig. 2a, right). This demonstrates that the distinct learning rules give rise to episodic and prototypical representations. To quantify these effects, we computed cosine similarity between DG representations and input images (Fig. 2b). DG_SUP_ representations exhibit significantly higher similarity to the original inputs than DG_INF_. Conversely, DG_INF_ representations are closer to class-average patterns than DG_SUP_ representations (Fig. 2c). Together, these results indicate that DG_SUP_ preferentially supports pattern separation, whereas DG_INF_ preferentially supports pattern integration.

**Figure 2.**
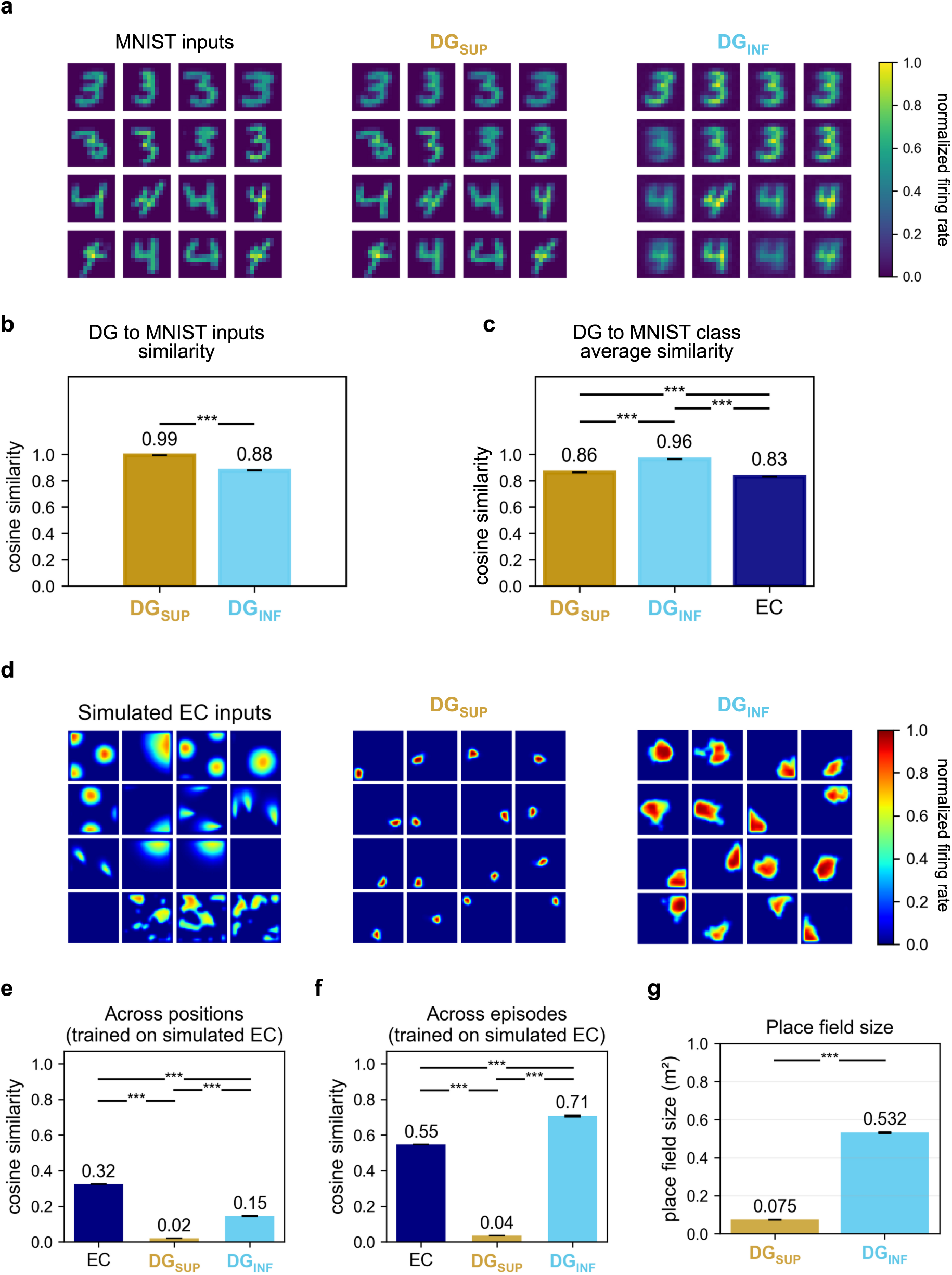
Separation and integration in DG representations. **a:** Sixteen randomly selected MNIST digits from classes 3 and 4 used as EC inputs, together with the corresponding DG_SUP_ and DG_INF_ reconstructions. DG representations were reconstructed as the activity-weighted average of receptive fields of active neurons. **b:** Cosine similarity between reconstructed DG representations and their corresponding input images, averaged across 500 MNIST images. DG_SUP_: 0.993, SD < 0.001; DG_INF_: 0.878, SD = 0.001. **c:** Cosine similarity between reconstructed representations and MNIST class-mean images. EC: 0.831; DG_SUP_: 0.863, SD < 0.001; DG_INF_: 0.964, SD = 0.001. **d:** Example positional tuning maps for 16 randomly selected neurons in simulated EC, DG_SUP_, and DG_INF_, shown on a normalized firing-rate scale. **e:** Cosine similarity between different positions within the same environment. EC: 0.324; DG_SUP_: 0.021, SD < 0.001; DG_INF_: 0.145, SD = 0.001. **f:** Cosine similarity between corresponding positions across different episodes within the same environment. EC: 0.548; DG_SUP_: 0.035, SD < 0.001; DG_INF_: 0.707, SD = 0.005. **g:** Mean place field size after segmentation. DG_SUP_: 0.075, SD < 0.001 m^2^; DG_INF_: 0.532, SD = 0.003 m^2^. Model values are reported as means with SDs across 50 independently seeded training runs per blade; EC values are fixed input-derived baselines. DG_SUP_–DG_INF_ comparisons used two-sided Welch independent-samples t-tests, whereas comparisons against the corresponding fixed EC baseline used two-sided one-sample Wilcoxon signed-rank tests. Holm correction was applied within each prespecified family of comparisons. All comparisons were significant after Holm correction. ***Adjusted p < 0.001.

We next tested whether these differences also apply to spatial inputs. Using the entorhinal cortex (EC) dataset, both networks were trained over five episodes in a square arena, in which object positions were slightly displaced. To visualize spatial selectivity, we plotted positional tuning maps of 16 randomly selected neurons for each blade (Fig. 2d). DG_SUP_ neurons exhibit sharper and more localized place fields (Fig. 2d middle), whereas DG_INF_ neurons display broader fields (Fig. 2d right). All positional tunings for both subnetworks are plotted in Fig. S3a and S3b. We further quantified place-field size using the peak-normalized and absolute thresholding criterion described in Methods, revealing significantly smaller fields in DG_SUP_ compared to DG_INF_ (Fig. 2g). We then examined positional decorrelation within episodes by computing pairwise similarities of population activity across spatial locations. Both DG blades show lower positional similarity than EC inputs, indicating increased separability across space, although DG_INF_ retains higher correlations due to broader fields (Fig. 2e). Thus, both blades disambiguate spatial positions, while DG_SUP_ shows the stronger pattern-separating regime. To assess representations across episodes, we computed cosine similarity between activity vectors at corresponding spatial locations across all episode pairs. DG_SUP_ representations are nearly completely decorrelated across episodes, indicating strong episodic separation (Fig. 2f). In contrast, DG_INF_ shows elevated cross-episode similarity, exceeding that of EC inputs, consistent with integration of information across episodes (Fig. 2f).

Together, these results demonstrate a functional dissociation between the DG blades, with DG_SUP_ implementing near-maximal pattern separation and DG_INF_ integrating information across episodes while preserving spatial structure.

### Remapping and stability of the place fields

Recent DG experiments have examined the stability and remapping of DG activity across repeated exposures to familiar and novel environments using linear tracks (Hainmueller and Bartos, 2018). To compare with these experiments, we generated a discrete trajectory through the same 2D environment and compared activity across different episodes of that environment. After training the model on all locations, we recorded the neuronal activity at each step along the linear trajectory for a single episode. For each comparison, neurons were retained if their maximum activity along the trajectory exceeded 0.05 in either episode. Among these neurons, those exceeding 0.05 in the reference episode were ordered by the position of their maximum activity. Neurons active above threshold only in the comparison episode retained their relative row positions within the filtered set. The resulting ordering was then applied unchanged to the second episode generated from the same environment distribution: geometry and object identities were preserved, but object positions were slightly perturbed and random spatial cell activity was resampled. We observed more pronounced remapping of place cells in DG_SUP_ (Fig. 3a, right) and largely stable place fields in DG_INF_ (Fig. 3b, right). For comparison, Hainmueller and Bartos (2018) show experimental data from mice navigating a linear track, where neurons are also sorted by peak firing location in one session, and the same ordering is applied to subsequent runs in a familiar environment and in a distinct environment. These in vivo recordings show a mixture of stable and remapping fields. This mixture of cells could potentially be explained by the coexistence of neurons with distinct coding regimes – neurons that integrate information across multiple similar patterns and hence fire for each of these patterns (captured by DG_INF_ in the model) and episode-specific neurons with strong pattern separation (captured by DG_SUP_). The extent to which these coding regimes are anatomically segregated remains an open question (cf. Fig. 7).

**Figure 3.**
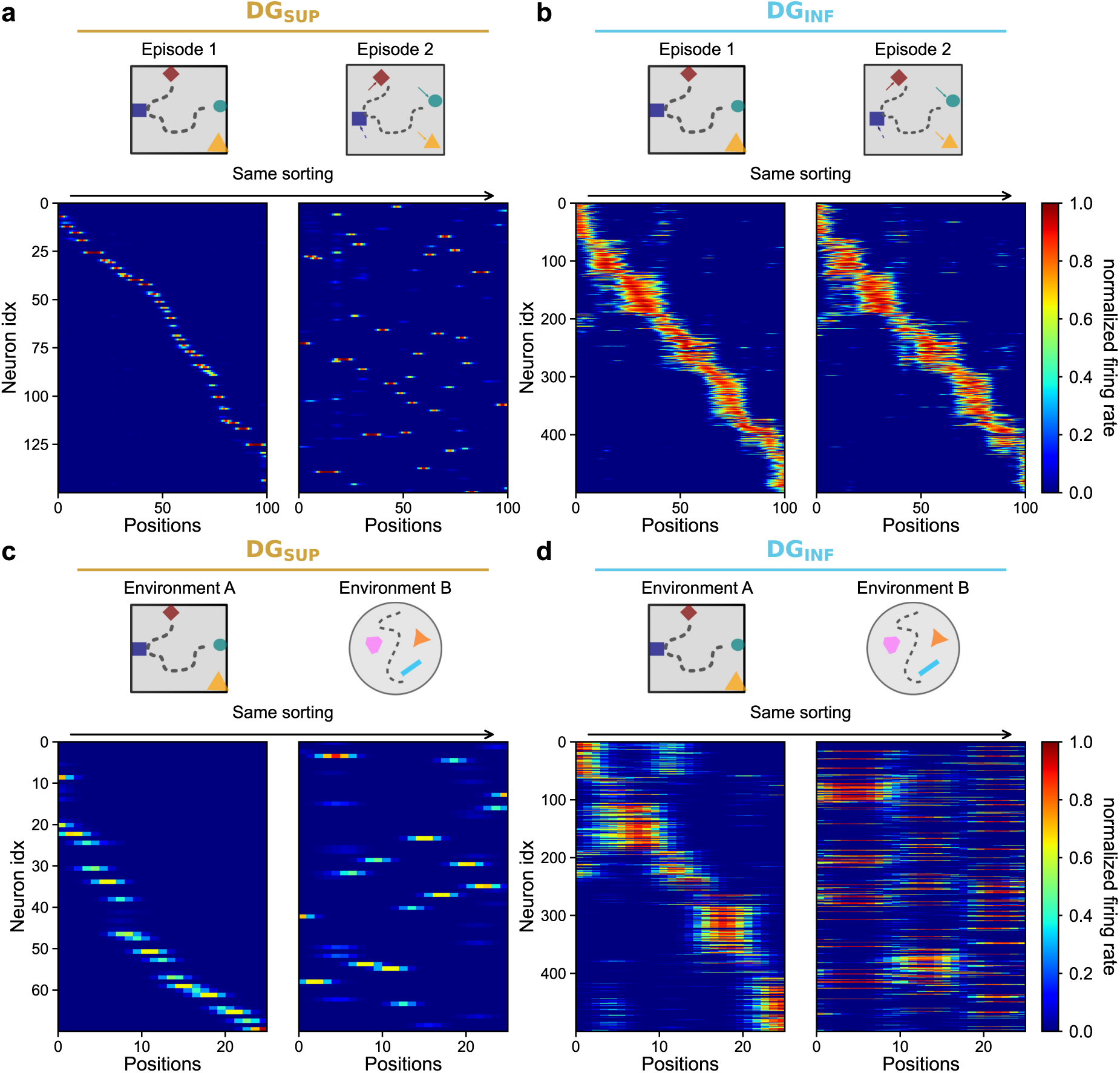
Remapping and stability of the place fields along a linear trajectory. **a–b:** Model DG activity along a linearized trajectory through the same 2D environment in two episodes. Neurons are sorted by their peak firing position in episode 1, and the same ordering is applied to episode 2, in which object positions are slightly perturbed and random spatial cell activity is resampled. DG_SUP_ neurons show pronounced episode-specific remapping (a), whereas DG_INF_ neurons preserve a largely stable diagonal structure across episodes (b), consistent with integration across similar experiences. **c-d:** Model DG activity along trajectories in two distinct environments: a square arena with four objects, environment A, and a circular arena with three different objects, environment B. Neurons are sorted by peak firing position in environment A, and the same ordering is applied to environment B. Both DG_SUP_ (c) and DG_INF_ (d) lose the diagonal structure across environments, indicating global remapping when geometry and object configuration change substantially. For c–d, inputs were subsampled to a 10 × 10 spatial grid to allow training over episodes in both environments (twice as many episodes as in a–b), yielding shorter linearized trajectories. In all model panels, neurons were retained if their maximum activity along the trajectory exceeded 0.05 in at least one of the two compared conditions. Among the retained neurons, those exceeding the threshold in the reference condition (episode 1 in a–b and environment A in c–d) were sorted by the position of maximum activity, and the resulting ordering was applied unchanged to the comparison condition.

Additionally, we construct a dataset with two environments with distinct geometry. Environment A is a square arena, which contains four objects. Environment B is a circular arena with three different objects. Simulated entorhinal inputs are modified accordingly: random spatial cell activity remaps, a different set of identity cells becomes active, and object-vector cell responses adjust to the new object locations. Grid-cell maps were assigned fixed environment-specific orientations and phase shifts in environment B, consistent with experimentally observed grid-cell remapping across environments (Fyhn et al., 2007; Stensola et al., 2012). Networks were trained on five episodes in each environment using a subsampled 10 × 10 grid, retaining the 88 positions shared by the square and circular environments (see Methods). In Fig. 3c, place fields are again sorted by the location of their peak firing along the trajectory, and the same neuron ordering is applied across environments. Under these conditions, DG place fields lose their diagonal structure and exhibit global remapping in both DG_SUP_ (Fig. 3c) and DG_INF_ (Fig. 3d). This contrasts with the stability of DG_INF_ observed across different episodes within the same environment (Fig. 3b). To control for within-environment variability, we additionally compared place fields across episodes within each environment, separately for the square and circular arenas. DG_SUP_ showed remapping across episodes, whereas DG_INF_ representations remained stable within each environment, as expected (Fig. S5).

Together, these results link blade-specific coding regimes and learning rates to experimentally observed mixtures of stable and remapping place fields and provide a mechanistic explanation for heterogeneous DG dynamics. Our model also predicts that when inputs correspond to distinct environments, for example because geometry and/or object identities change substantially, DG_INF_ no longer integrates them into a common representation, but instead forms separate integrated maps for each context, at least as far as spatial coding is concerned.

### Separability of representations across positions and episodes

We computed pairwise separability to quantify how separated patterns are across all activity patterns, but also across positions or episodes (Fig. 4a). When trained on MNIST inputs, both DG subnetworks increase the separability metric relative to EC inputs, with DG_SUP_ approaching near-maximal separability (Fig. 4b). This effect persists also when we consider the separability only within the same class (Fig. 4d). Here, we note that the separability increases for both DG_SUP_ and DG_INF_ representations compared to EC due to the increased size of the DG layer when the model is trained with MNIST. This leads to an increase in separability, albeit the representations remain close to the averages in DG_INF_ and to exemplars in DG_SUP_ (see Fig. 2b and Fig. 2c). When we analyzed separability after training on simulated neural EC activity, DG_SUP_ again approached maximal separation. DG_INF_ showed an intermediate regime: separability was substantially lower than in DG_SUP_, but higher than in EC when considering all spatial activity patterns (Fig. 4c). A similar pattern was observed for different positions within the same episode, where DG_INF_ increased positional separability relative to EC but remained less separated than DG_SUP_ (Fig. 4e). The distinctive signature of DG_INF_ emerged when comparing activity patterns corresponding to the same spatial location across different episodes: here, DG_INF_ separability was lower than that of the EC inputs (Fig. 4f), indicating cross-episode integration. In contrast, DG_SUP_ maintained high separability across both positions and episodes.

**Figure 4.**
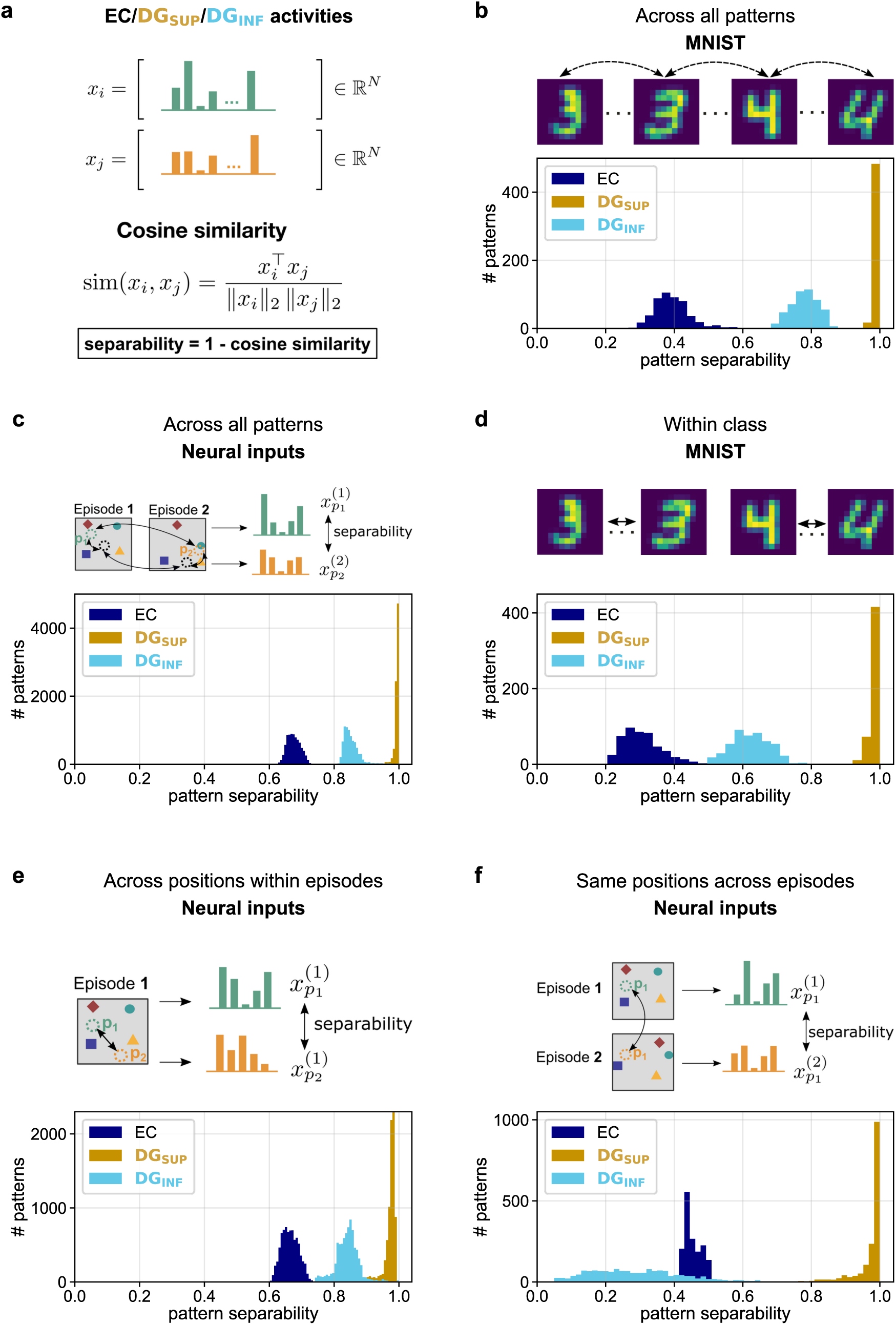
Separability of representations in EC, DG_SUP_, and DG_INF_ for MNIST and simulated neural data. **a:** Schematic of the separability analysis. For each activity pattern, separability was defined as 1 minus the mean cosine similarity to the other activity patterns in the relevant comparison set. **b:** Separability across all MNIST patterns (patterns from classes 3 and 4). EC: 0.394 ± 0.053; DG_SUP_: 0.993 ± 0.007; DG_INF_: 0.780 ± 0.040. **c:** Each pattern was compared with all other simulated patterns, including patterns from different positions within the same episode, the same position across different episodes, and different positions across different episodes. *x*^(*e*)^ - activity at position *p_i_* in the episode *e*. EC: 0.676 ± 0.020; DG_SUP_: 0.993 ± 0.006; DG_INF_: 0.854 ± 0.027. **d:** Separability among MNIST patterns belonging to the same class, pooled across classes 3 and 4. EC: 0.311 ± 0.063; DG_SUP_: 0.986 ± 0.015; DG_INF_: 0.624 ± 0.057. **e:** Separability between activity patterns representing only different positions within the same episode. The separability values were pooled across all episodes. EC: 0.664 ± 0.026; DG_SUP_: 0.973 ± 0.019; DG_INF_: 0.841 ± 0.036. **f:** Separability between activity patterns representing only the same spatial position across different episodes. The separability values are then averaged across all positions. EC: 0.454 ± 0.025; DG_SUP_: 0.970 ± 0.047; DG_INF_: 0.291 ± 0.133. Values are mean ± SD across patterns (**b, d**: n = 500; c: n = 8,000), episode-position observations (**e**: n = 8,000), or spatial positions (**f**: n = 1,600). These distributions were computed from one trained model per blade and dataset rather than from activities pooled or averaged across independently seeded runs. Accordingly, these SDs describe variability across the stated within-model analysis units and do not quantify variability across model initializations.

Overall, DG_SUP_ consistently amplifies differences between input patterns, consistent with a strongly pattern-separating regime. By contrast, DG_INF_ occupies an intermediate regime: it preserves and increases separability across spatial positions relative to EC, but reduces separability between corresponding positions across episodes. This pattern is consistent with integration across related experiences without collapse of the underlying spatial structure.

### Functionally combining blades for novel memory encoding and forgetting

To link DG representations to downstream function, we trained linear decoders to reconstruct entorhinal cortex (EC) inputs from DG activity for each blade and dataset. Decoders were trained on the same patterns and episodes used to train the DG_SUP_ and DG_INF_ networks. Prediction error was operationalized as decoder reconstruction error, computed from the cosine similarity between the decoded EC pattern and the corresponding true EC input. High cosine similarity indicates that the current input is well predicted by the blade-specific DG representation, whereas low cosine similarity indicates a large prediction error (Fig. 5a). This provides a signal that quantifies how well each blade predicts ongoing input. Crucially, this justifies the coexistence of separation and integration of memories beyond a visual match to experimental recordings. That is, we can derive a concrete functional reason for combining integration and separation streams: comparing blade-specific signals can guide encoding of novel patterns and forgetting of well-represented memories. We hypothesize that the blade-specific prediction errors could be read out separately and compared with the corresponding EC input. Four cases of high or low prediction errors in one or both of the blades can occur (Fig. 5d):

- **Low error in both blades** indicates that the current input is well predicted by an existing prototype in DG_INF_ and that a similar episodic trace exists in DG_SUP_. In this regime, the episodic trace is redundant with the learned integrated representation. Hence neuronal resources can be freed up. For instance, without continued reinforcement, such traces may weaken over time, consistent with synaptic tagging and capture mechanisms (Barco et al., 2008; Frey and Frey, 2008) and neurogenesis-associated turnover (Akers et al., 2014).
- **Low error in DG_INF_ but high error in DG_SUP_** indicates that the input matches an existing prototype (formed across several experiences) whereas the corresponding episodic representation has already been removed from DG_SUP_ by the forgetting process. In this case, no novel encoding or forgetting is needed.
- **High error in both blades** signals a novel input that is not captured by existing prototypes or exemplars. This requires the formation of a new episodic memory in DG_SUP_ and also gradual integration of the memory into a new prototype in DG_INF_.
- **Low error in DG_SUP_ but high error in DG_INF_** indicates that an episodic memory exists, but there is no corresponding prototype. In this case, the prototype should be gradually formed in DG_INF_.

**Figure 5.**
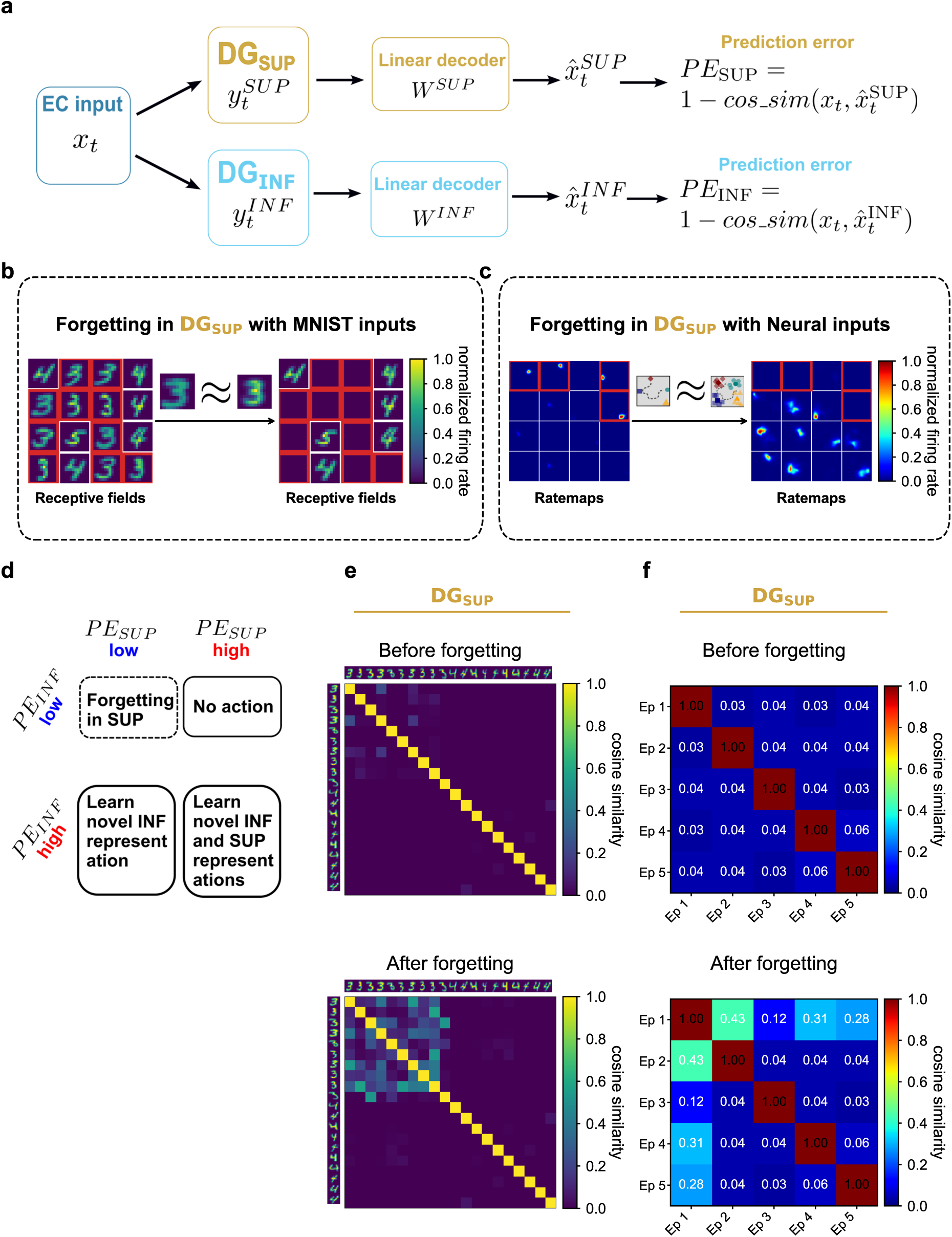
Decoding and prediction-error–guided forgetting in DG_SUP_. **a:** Schematic of the decoding procedure used to compute blade-specific prediction errors. EC input activity is reconstructed from DG_SUP_ and DG_INF_ activity using linear decoders, yielding separate PE_SUP_ and PE_INF_ signals. **b:** Forgetting in DG_SUP_ for MNIST inputs. When an input is well predicted by DG_INF_ and a corresponding DG_SUP_ trace is still detectable, the episodic representation is treated as a candidate for deletion and reset in DG_SUP_. Receptive fields are shown before and after forgetting; neurons selected for forgetting are highlighted in red. **c:** Forgetting in DG_SUP_ for simulated neural inputs. Positional tuning maps are shown before and after forgetting. Affected neurons lose spatial selectivity after the corresponding DG_SUP_ units are reset. **d:** Summary of the four possible prediction-error regimes and their proposed functional consequences: forgetting, no action, integration in DG_INF_, or encoding in both blades. Panels b and c show the effects of forgetting on receptive fields and positional tuning, whereas panels e and f show the resulting changes in representational similarity. **e:** Pairwise cosine similarity between DG_SUP_ representations of selected MNIST digit-3 and digit-4 exemplars before and after forgetting. Forgetting candidates were selected from the digit-3 exemplars. Before forgetting, each exemplar preferentially activates a dedicated neuron, producing distinct population representations. Forgetting deletes and resets selected exemplar units. Consequently, the corresponding inputs are represented by remaining committed neurons encoding similar exemplars, thereby increasing representational overlap and pairwise similarity. **f:** Pairwise cosine similarity between DG_SUP_ representations before and after threshold-gated forgetting applied to candidate patterns from one trained simulated EC episode. Similarity between episode representations increases after forgetting.

We first demonstrate prediction-error–guided forgetting for MNIST inputs (Fig. 5b). A selected instance of the digit “3” lies close to the learned DG_INF_ prototype and is therefore well reconstructed from the integrated representation. When a corresponding DG_SUP_ trace is still detectable, this indicates that the same input is represented both as an integrated prototype-like code and as an episodic exemplar. Such traces are treated as candidates for forgetting, implemented as deletion and reset of the corresponding active DG_SUP_ units (see Methods). The same principle applies to the DG_SUP_ subnetwork driven by simulated EC inputs, where neurons undergoing forgetting show corresponding changes in positional tuning (Fig. 5c). We also show how representational similarity of the DG_SUP_ patterns changes after DG_SUP_ units associated with multiple candidate patterns have been deleted and reset when trained on the MNIST dataset (Fig. 5e) and simulated EC inputs (Fig. 5f). Specifically, pairwise representational similarity increased after forgetting, corresponding to reduced pattern separation.

Next, we examine how prediction errors can drive the formation of new representations in both subnetworks. We propose a mechanism, in which encoding of novel patterns in both blades is supported by neurogenesis-inspired turnover/recruitment. In our model, it is implemented as a turnover process, in which neurons with weak connectivity are re-initialized and selected for encoding. We then suggest that the GABA-switch mechanism of maturation of adult-born granule cells can support the formation of novel integrated prototypes in DG_INF_ (Fig. 6d). In the early phase before the switch, elevated intracellular chloride makes GABAergic interneuron input excitatory, promoting cooperative activity and broad tuning. In the late phase, GABAergic input switches to inhibitory due to the decrease in the intracellular chloride concentration. This switch to inhibition enables competitive refinement of selectivity (Ge et al., 2006; Gozel and Gerstner, 2021). Consistent with this mechanism, newly recruited neurons (Fig. 6a, left) initially exhibit broad, prototype-like responses to a novel MNIST class (Fig. 6a, middle), before developing more selective, but still integrated, class-level representations in the late phase (Fig. 6a, right). A similar progression is observed when a novel same-environment EC episode is encoded in the network trained with simulated EC inputs: neurons are broadly active across all positions during the early phase (Fig. 6b, middle) and develop distinct place fields after maturation, while mature neurons that are not selected for turnover either preserve their positional tuning or change only slightly (Fig. 6b, right). Hence, encoding of a novel same-environment episode can be supported through neurogenesis-inspired turnover and a GABA-switch-like maturation mechanism during refinement of the existing prototype. By contrast, when a novel distinct-environment episode is encoded, newly recruited neurons acquire novel positional tunings for the new environment, while mature neurons largely remain silent (Fig. 6c). This constitutes the initial formation of a separate integrated representation for the new environment. For DG_SUP_, we propose a similar mechanism involving the turnover of weakly connected neurons for novel memory encoding. Here, an explicit GABA-switch mechanism is not necessary, as the neurons do not need to undergo the cooperative phase. Hence, we suggest that novel inputs are encoded via recruitment of new adult-born granule cells, resulting in rapid formation of exemplar representations. We observe that recruited weakly connected neurons with no previous receptive fields (Fig. 6e, left) are able to form exemplar representations for a novel MNIST class in DG_SUP_ after one-shot learning (Fig. 6e, right). The same process occurs for simulated EC inputs: neurons form spatially selective place fields both for the novel same-environment episode (Fig. 6f) and for the novel distinct-environment episode (Fig. 6g). Unlike DG_INF_, this process does not include an explicit GABA-switch mechanism due to the absence of modeled interneurons, and therefore captures only the fast encoding regime. Correspondingly, similarity structure diverges across the two blades: DG_SUP_ representations become more decorrelated after encoding, whereas DG_INF_ representations show full integration during the early phase, followed by the stabilization of more refined integration after maturation (Figs. S6-S8).

**Figure 6.**
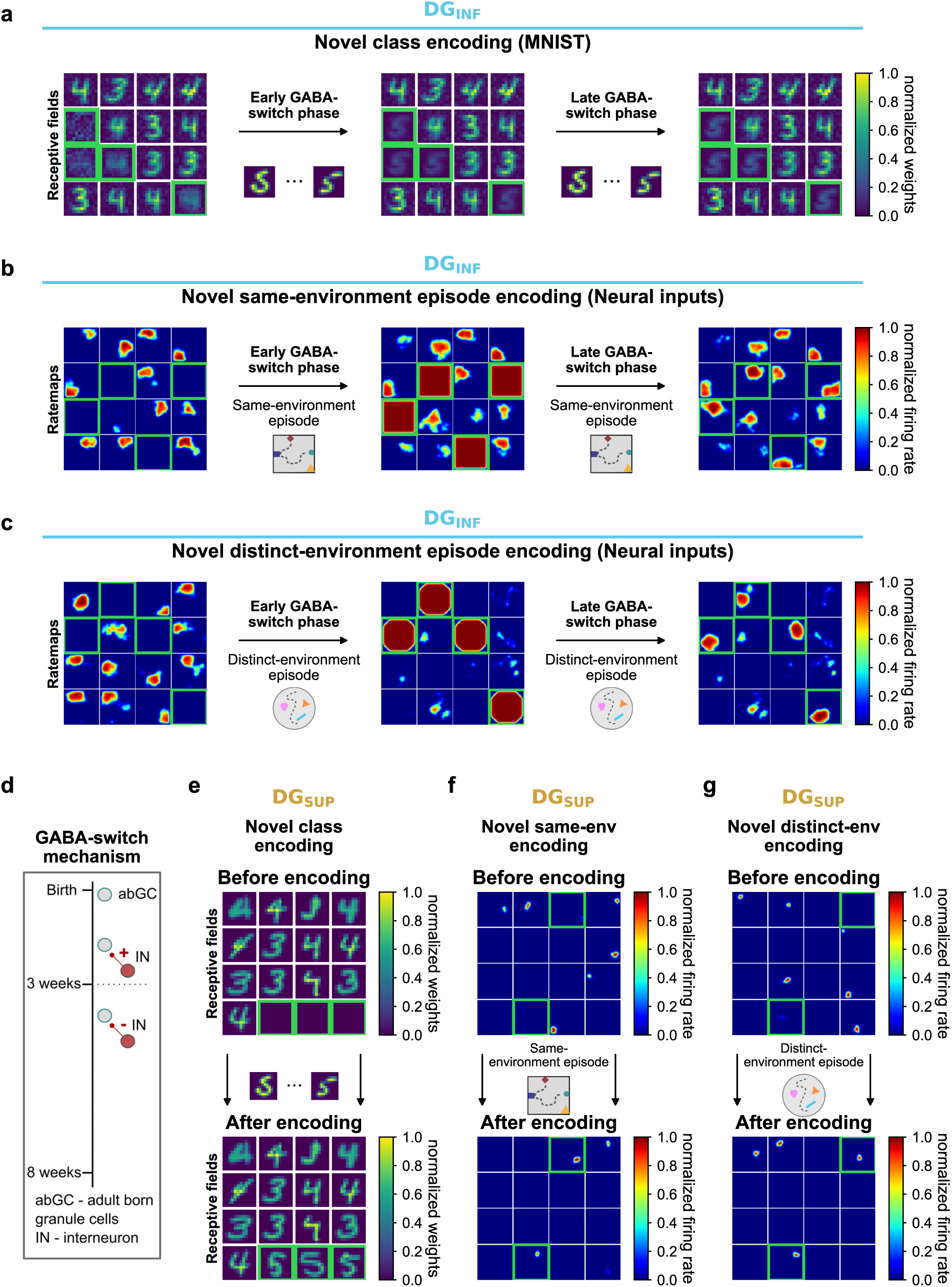
Novel memory encoding in DG_INF_ and DG_SUP_. **a:** Novel-class encoding in DG_INF_ for MNIST inputs. Receptive fields are shown before encoding, after the early GABA-switch phase, and after the late GABA-switch phase. Newly recruited neurons are highlighted in green. Early excitation produces broad receptive-field responses, whereas the late inhibitory phase refines selectivity. **b:** Novel encoding of a same-environment episode in DG_INF_ for simulated neural inputs. Positional tuning maps are shown before encoding, after the early phase, and after the late phase. Newly recruited neurons initially respond broadly due to excitation from inhibitory cells and later develop more localized place fields because of the switch to inhibition. **c:** Novel encoding of a distinct-environment episode in DG_INF_. As in **b**, early recruitment produces broad responses, followed by the formation of a distinct integrated representation after the late phase. **d:** Schematic of the GABA-switch in adult-born granule cells, showing the transition from early depolarizing to later hyperpolarizing GABAergic input during maturation. **e:** Novel-class encoding in DG_SUP_ for MNIST inputs. Weakly connected neurons are recruited and acquire receptive fields for novel digit-class exemplars after one-shot learning. **f–g:** Novel encoding in DG_SUP_ for simulated neural inputs. Recruited neurons form new place fields after one-shot encoding of either a novel same-environment episode **(f)** or a novel distinct-environment episode **(g)**.

## Discussion

This study constitutes the first computational model that differentiates between the two blades of the dentate gyrus, assigns them specific computational roles, and explores how these two streams can be functionally combined. This has implications for differences in downstream processing since the two blades preferentially target different parts along the proximo-distal extent of CA3 (Witter, 2007a). The model also goes beyond the hitherto dominant proposal of pure pattern separation in the DG. It consists of two distinct networks: EC-DG_SUP_ and EC-DG_INF_. Each network is trained using a different learning rule: a k-WTA-based learning mechanism with rapid recruitment and one-shot imprinting (high learning rate) for DG_SUP_ and a heterosynaptic rule with a small learning rate for DG_INF_. We show that the two blades learn distinct representations: more rapidly acquired episodic representations in the DG_SUP_ and slowly accumulating representations in the DG_INF_. This interpretation is consistent with recent experimental work showing that DG population codes refine gradually over repeated exposures, with increasing sparsity, specificity, and reliability over days (Cholvin and Bartos, 2026). Here “slow” should be put in context. The DG_INF_ model does not require an unrealistically high number of training events (five generated episodes with a few repeated training passes are sufficient), despite not implementing one-shot learning, nor does it employ biologically implausible learning rules. Driven by EC-like inputs and the distinct learning dynamics of each blade, place fields emerge in both DG blade networks. We show how the model may account for the stable firing of DG cells across days, and propose that the two distinct coding regimes are used to gradually build cognitive maps while preserving individual episodes when necessary (see discussion of prediction errors below).

The k-WTA mechanism in the suprapyramidal blade selectively allocates neurons to inputs based on a similarity threshold, resulting in narrowly tuned, highly specific representations compared to the broader fields observed in the infrapyramidal blade. Because recruitment is triggered when inputs are insufficiently similar to existing representations, this mechanism provides a simple account of how new place-cell-like responses could be formed as an agent moves through space. We speculate that such a drop in pattern similarity as an agent moves in space could constitute a valid recruitment mechanism for place cells in general (including in downstream CA3), and that the suprapyramidal-infrapyramidal blade distinction carries over to proximal vs distal CA subfields in line with Berdugo-Vega et al. (2023). The bimodal distribution of place field size in DG_INF_ vs DG_SUP_ could be detectable in DG recordings, though the possibility remains that this difference is attenuated by factors not modeled here or that the anatomical distribution is not as strict (see below). Broader place fields may potentially facilitate the integration capabilities of DG_INF_ neurons. Population vector (PV) analysis suggests that both DG blades preserve positional separability, as indicated by reduced position similarity relative to EC inputs. However, their dynamics across episodes diverge: DG_SUP_ shows decreasing similarity across episodes, consistent with episodic separation, whereas DG_INF_ shows increasing similarity across episodes, consistent with integration while preserving positional structure. This functional dissociation is reinforced by remapping in DG_SUP_ and stable representations in DG_INF_ across episodes from the same environment distribution, supporting a dual role for the DG in both separating and integrating episodic content. Recent work in CA1 showed the coexistence of the stable and unstable place cell populations across several days (Climer et al., 2025). Although intrinsic membrane excitability was not measured directly, cell stability was linked to functional measures of excitability, including average ΔF/F, calcium-transient integral, and the percentage of active laps, which together predicted subsequent stability. These findings additionally support the broader principle that stable and unstable/plastic coding regimes may coexist across the hippocampal circuit.

Our model lives at the algorithmic level of Marr’s theoretical framework (Marr, 1982), albeit with many references to detailed biological constraints, such as plausible connection motifs and a modest number of learning epochs that could be accounted for by neural replay. The suprapyramidal blade is implemented as a feedforward competitive network without explicit continuous dynamics, whereas the infrapyramidal blade includes rate-based excitatory and inhibitory dynamics. Future extensions of the model could investigate more neural details through the incorporation of the diverse cell types found in the dentate gyrus, and address differential contributions of LEC and MEC to the two blades (Vivar et al., 2012; Woods et al., 2018). Guided by the computational hypotheses proposed here, this could contribute to a better understanding of the roles of individual cell types in the suprapyramidal and infrapyramidal blades. Furthermore, we focused mainly on modeling the DG networks without explicitly modeling the downstream regions such as CA3, CA1, or the subiculum.

Across both MNIST and simulated neural datasets, our analysis of representational geometry shows that DG_SUP_ strongly decorrelates activity patterns, consistent with episodic pattern separation. Conversely, DG_INF_ maintains higher similarity among related inputs: it preserves high separability across spatial positions, but shows reduced separability for corresponding positions across episodes. Thus, the model captures key aspects of DG function: spatial pattern separation across positions, episodic separation in DG_SUP_, and gradual cross-episode integration in DG_INF_.

Having both exemplars (episodes) and averages (prototypes, as a result of integration across episodes) at its disposal, we suggest that DG can underlie prediction errors that can guide the formation and forgetting of episodic memories. That is, DG could inform CA1 and EC, previously suggested to be involved in prediction error calculations (Bein et al., 2020; Ku et al., 2021). We also propose that prediction errors could similarly be computed individually for each blade. Combined prediction errors from both blades could inform whether episodic memories have been sufficiently integrated into generalized representations, thereby enabling synaptic repurposing during neuronal turnover. High prediction errors across both blades indicate a need for new episodic and integrated representations, whereas low prediction errors indicate that the input is already well captured by existing representations. We further illustrate how novel memory encoding may be supported through the recruitment of adult-born granule cells, utilizing heterosynaptic plasticity and the GABA-switch mechanism (Ge et al., 2006), implicitly introducing aspects of turnover/neurogenesis into the model. In early phases, recruited neurons cooperate broadly, while late-phase neurons compete to establish specific memory selectivity, consistent with previous work (Gozel and Gerstner, 2021). We demonstrate learning of new memories using both the MNIST dataset (with a new digit class) and a sufficiently distinct environment for the neural input dataset.

Finally, a prior theoretical framework provides additional motivation for the notion of comparing predictions to current experience. The SPEAR model has suggested that the theta rhythm determines encoding vs retrieval phases in the hippocampus proper (Hasselmo et al., 2002). This would allow for memory readout (CA3 to CA1) without interfering with the establishment of novel associations within CA3 and between CA3 and CA1. However, if memory readout is orchestrated at the frequency of theta (about 8 times a second), this cannot entail a full cascade of pattern completion from the hippocampus to various cortical areas as must occur during conscious recall (Staresina and Wimber, 2019). Instead, this “memory-readout” must be different from full recollection, and could potentially be more anatomically localized. We propose that these theta dynamics, consistent with the earlier proposal by Hasselmo and colleagues, could allow for a comparison - at the level of the hippocampus - of generalized expectations and current experience, in addition to any interference-mitigating aspects. That is, during ongoing behavior, current experience can be checked against expectations about what to find at a given location. In the present model, this corresponds to a particular object configuration. More generally, we can liken extensive coverage of an environment with well-developed expectations to a cognitive map formed from overlapping episodes.

We suggest that integration should occur primarily across episodes in the same environment, with less regard to temporal proximity of experiences across environments. Nevertheless, this does not necessarily exclude integration across time for disparate events close in time. However, how two very different experiences (in different environments) should be integrated when occurring in temporal proximity is beyond the scope here and would likely require additional machinery. Indeed, the temporally proximal events are potentially only integrated within the hippocampus when they share the same structure, integrating temporal relations across similar sequences (Bellmund et al., 2022). The utility of integrating temporally close experiences could lie in the establishment of temporal relations between disparate events, while integrating very similar experiences (even if days apart) can yield the kinds of averages (or expectations) we show here, useful for constructing a cognitive map from overlapping episodes in the same environment.

Very recent, direct experimental support for the model can be found in the study by Strauch et al. (2025), reporting blade-specific differences in synaptic plasticity in the dentate gyrus. The suprapyramidal blade exhibited notably stronger long-term potentiation (LTP) and long-term depression (LTD) compared to the infrapyramidal blade, a difference that coincided with lower GluN subunit expression in DG_INF_ relative to DG_SUP_. This is consistent with the role of NMDARs containing these subunits in facilitating LTP in the dentate gyrus (Vasuta et al., 2007). Previous computational models of the DG have used biologically plausible EC inputs (Aimone et al., 2009; Appleby et al., 2011; Cuneo et al., 2012; Finnegan et al., 2017). However, most of these inputs were relatively simplified and did not incorporate object-related cell types. In contrast, our model utilizes environments populated with multiple objects and EC neuron populations selective for object identity and location, thus better approximating episodic experiences.

In related work, Berdugo-Vega et al. (2021) demonstrated that increasing neurogenesis through stem-cell expansion enhanced DG_INF_ activity during spatial navigation tasks, resulting in more precise and directed trajectories after the animal learned the task. Conversely, reversal trials showed decreased DG_SUP_ activity, associated with reduced perseveration at previously rewarded locations. The precision of the trajectories may indicate the formation of integrated representations in the DG_INF_, whereas the reduced persistence of trajectories near the previously rewarded locations suggests improved contextual discrimination between the standard and reversed trials. These findings support the hypothesis of distinct pattern separation and integration streams, underpinned by preferential connectivity of LEC to DG_SUP_ and MEC to DG_INF_ and subsequent hippocampal layers (Berdugo-Vega et al., 2023; Tamamaki, 1997; Vivar et al., 2012; Witter, 2007b).

Additionally, the study by Luna et al. (2019) showed that the modulatory effect of the abGCs is dependent on the incoming inputs: abGCs with inputs from LEC inhibit mGCs, whereas abGCs with inputs from MEC excite mGCs. This implies greater inhibition in DG_SUP_ and greater excitation in DG_INF_, consistent with the proposed separation and integration streams.

In experiments with mice navigating linear virtual tracks, Hainmueller and Bartos (2018) observed stable DG place fields across multiple days, whereas CA3 and CA1 place fields differed significantly across days. Correlations between familiar and novel environments were notably higher in DG compared to CA3 and CA1. Hainmueller and Bartos (2018) also showed higher place field stability in the proximal CA3 compared to the distal CA3, which is compatible with the hypothesis that the integration and separation processing streams continue throughout the hippocampus. These findings indicate integration capabilities in at least a subset of DG place cells. In an analogous analysis, our model showed pronounced remapping of place fields in DG_SUP_ and relative stability in DG_INF_ across different episodes of the same environment. This could reflect recordings that combine cells from both blades, recordings of axons extending from one blade into the other, or a less strict anatomical separation. The anatomical distribution of neurons from our episodic and integration networks could in principle take many forms (cf. Fig. 7). There is some evidence for significant functional distinctions between blades (Luna et al., 2019; Berdugo-Vega et al., 2021, 2023; Strauch et al., 2025), but alternatively the two processing streams could show more overlap (Fig. 7b, c) or their relative proportion could change along the septo-temporal axis of the hippocampus. Future experiments can determine the validity of our computational proposal and (separately) its anatomical distribution. Similarly, there is evidence that abGCs actively remodel the place cells in CA3 (Mugnaini et al., 2023), while mature cells maintain stability or facilitate pattern completion (Nakashiba et al., 2012). These findings imply direct encoding by abGCs, not merely modulation of mGCs. Potential blade-specific differences in the recruitment and encoding roles of abGCs and mGCs seem promising targets for future investigation.

**Figure 7.**
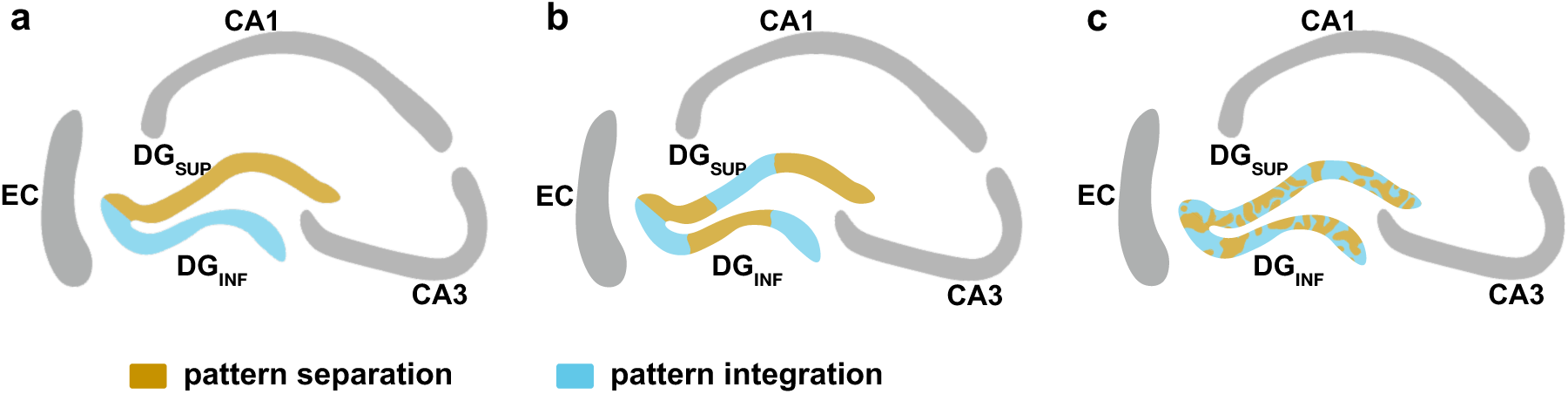
Possible anatomical distributions of episodic versus integrated processing in DG. **a:** Segregated organization: episodic (DG_SUP_-like) and integrated (DG_INF_-like) representations are largely confined to separate blades. **b, c:** Distributed organization: episodic and integrated representations coexist within both blades, with varying proportions or spatial clustering. Not shown: the relative contribution of episodic versus integrated representations may additionally vary along the septo-temporal axis of the hippocampus.

In summary, we present the first model of the DG to distinguish blade-specific computation and learning. It proposes how episodic-like one-shot learning and integration across experiences could be accomplished in the DG with biologically plausible learning regimes, and how these two streams can interact fruitfully. The combined system thus accomplishes more than the sum of its parts. The model makes several predictions, suggests how to account for previously unexplained experimental data, and proposes how the resultant representations could inform further hippocampal processing, painting the DG as a key player in the hippocampus, beyond its well-established role in pattern separation.

## Methods

### Model inputs

We use two datasets as inputs to the model: the MNIST handwritten digit dataset (Lecun et al., 1998) and a simulated neural dataset consisting of cell populations found in the entorhinal cortex. The MNIST dataset contains 70,000 examples of handwritten digits from 10 classes. For training, we use a subset of 500 examples from 2 classes. To reduce input dimensionality, each image is downsampled from 28 × 28 to 12 × 12pixels and flattened to an *N_EC_* = 144-dimensional vector.

To simulate entorhinal cortex inputs, we define several cell populations: grid cells (245 cells), object-vector cells (256 cells), identity cells (252 cells), and random spatial neurons (250 cells). We select approximately equal numbers of cells for each population, while ensuring complete positional coverage for grid cells and balanced coverage across objects for object-vector and identity cells. Each simulated input pattern therefore has dimensionality *N_EC_* = 1003. During each episode in the single-environment experiments (Figs. 2, 3a and 3b, 4, 5, 6), the agent visits all 40 × 40 uniformly distributed positions in a square 2 m × 2 m environment. For the two-geometry experiment (Fig. 3c and 3d), we use a subsampled 10 × 10grid and retain only positions that exist in both the square and circular geometries, resulting in 88 shared positions. The environment is discretized for simplicity of training, although in principle the agent could also be moved along a realistic trajectory, for example generated with RatInABox (George et al., 2024).

Object-vector cells and random spatial neurons were generated using RatInABox. Identity cells were implemented as an additional class in RatInABox. Grid-cell firing fields were adapted from a previous model (Bicanski and Burgess, 2019). Prior to training, each input pattern (population activity vector) was normalized to unit *L*_2_norm, ∥ *x* ∥_2_= 1.

### Grid cells

Grid cells are modeled as precomputed firing-rate maps generated from three superimposed cosine waves with orientations of 0°, 60°, and 120°:

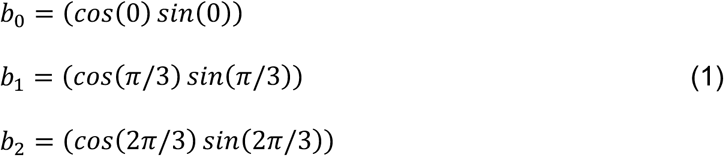

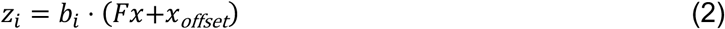

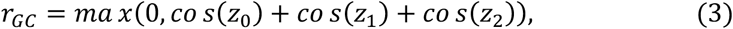

where b_0_, b_1_, and b_2_ are the basis vectors of the three cosine components. Frequencies *F* are defined separately for each module, starting from 0.0028 *x* 2*π* for the first module. Module scales follow experimental observations, such that successive modules differ by a factor of 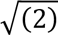 (Stensola et al., 2012). Modules also have different predefined orientations. Individual cells are offset relative to each other to ensure uniform spatial coverage. We use 5 modules with 49 cells per module.

### Object-vector cells

The RatInABox package was used to generate a population of allocentric vector-tuned neurons, with each cell responding to every object in the environment. First, a wall-aware vector, defined by distance and bearing from the query position to each object, is computed. A Gaussian tuning curve is applied over distance and a von Mises tuning curve over angle. The resulting responses are summed across all objects.

### Random spatial neurons

RatInABox was also used to generate a population of random spatial neurons. In RatInABox, these are implemented as a non-parametric Gaussian-process model of spatially tuned firing fields. For each neuron, a smooth random function is sampled from a Gaussian process with mean 0 and squared-exponential kernel with spatial length scale *l* = 0.1 m. A sigmoid transformation is applied to keep the firing rate within the range [*fr_min_*, *fr_max_*]. To compute firing rates, the covariance between query points and discretized locations is evaluated, kernel weights are normalized, and a weighted sum over the sampled function values is obtained. RSN activity values below 0.6 were set to zero before the different cell populations were combined into EC input vectors.

### Identity cells

Identity cells are modeled as Gaussian-tuned responses centered on object locations, with spatial standard deviation of 0.4 m.

### Different episodes in the same environment

Different episodes within the square environment were generated by varying the positions of four objects. One object-cluster center was sampled once from a predefined region in each quadrant and then held fixed across episodes. Episode-specific positions were sampled from object-specific bivariate Gaussian distributions. The x/y variances were 0.07/0.03, 0.10/0.08, 0.02/0.15, and 0.05/0.07 m², with x–y covariances of 0, 0.03, 0, and −0.02 m², respectively. Grid-cell activity remained unchanged, object-vector- and identity-cell responses followed the sampled object positions, and random spatial neurons were reinitialized for each episode.

### Different environments

Distinct environments differed in geometry, object configuration, and the activity patterns of all four EC populations. Environment A was a square arena containing four objects, whereas environment B was a circular arena containing three different objects. Random spatial neurons were independently reinitialized for every episode in both environments, producing episode-specific random spatial fields. Non-overlapping identity-cell subsets represented the objects in the two environments: identity slots 0–3 were active in environment A and slots 4– 6 in environment B. Object-vector-cell tuning parameters were held fixed, but their responses were recomputed from the environment-specific object positions and therefore changed between environments. Grid-cell maps were unchanged in environment A. Environment B did not use a single common rotation; instead, each grid module was assigned a fixed environment-specific orientation and phase shift. For the five grid-cell modules, orientations were [*π*/3, *π*/4, *π*/2, *π*/6,1.2*π*] radians in environment A and [0.10, 0.30, 0.213, 0.547, 0.062] radians in environment B. In the two-environment simulations, environment A was a square arena containing four objects sampled from quadrant-specific distributions, whereas environment B was a circular arena containing three objects arranged around predefined positions. In environment B, object positions varied between episodes using Gaussian positional jitter with SD 0.16 m, were constrained to remain inside the arena, and targeted a minimum separation of 0.9 m.

### Network architecture and learning The suprapyramidal blade network

The suprapyramidal blade network consists of an input layer (EC population activity) and a dentate gyrus layer. The model does not simulate continuous rate dynamics; instead, learning uses k-Winner-Take-All (k-WTA) winner selection with learned lateral inhibition, followed by a soft similarity-based activity readout. For each input pattern, DG neurons compute a similarity score

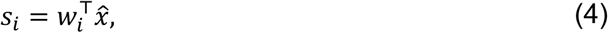

where *w_i_* denotes the feedforward weight vector of neuron *i* and *x*” is the normalized input. At input presentation *t* only neurons belonging to the pre-existing committed set *C_t_* participate in similarity matching and k-WTA competition. Novelty is determined by the maximal similarity among these neurons:

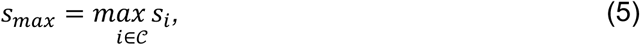

If *s_max_* < *θ_SUP_,* the input is considered novel and a previously unrecruited DG neuron is recruited. Recruitment implements competitive allocation of representational resources in response to novel inputs. Newly recruited neurons undergo one-shot Hebbian imprinting:

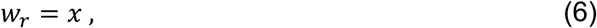

thereby storing the current input as an exemplar representation. The recruited neuron is added to the committed set for the next input presentation, *C_t_*_+1_ = *C_t_* ∪ {*r*}. Recruitment was treated as an immediate storage operation but with delayed activity eligibility. If no neurons were committed at the first presentation (*C_t_* = ∅), the input was necessarily classified as novel, a free neuron was selected at random and imprinted, and competition began from the subsequent presentation. It therefore becomes eligible for similarity matching, k-WTA competition, lateral-inhibition updates, and activity readout from the subsequent input presentation onward. Competition among neurons eligible at the current presentation is performed via k-WTA with learned lateral inhibition. This step selects winner neurons for exemplar reuse and lateral-inhibition updates. DG output activity is computed using a soft similarity-to-rate transformation over the current eligible set *C_t_*:

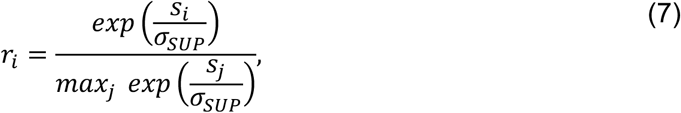

restricted to committed neurons. For winner neurons with (*s_i_* ≥ *θ*_SUP_), the weights of the winner neurons are additionally updated according to the following rule:

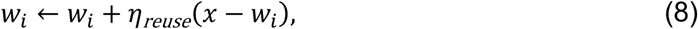

enabling limited integration of near-identical patterns. To reduce redundancy, the model learns an effective lateral inhibition matrix *L*. After winner selection, inhibition from winner *i* to competitor *j* is strengthened proportionally to competitor similarity:

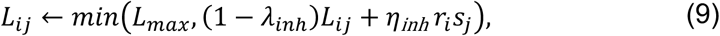

with decay *λ_inh_* and inhibitory learning rate *η_inh_*. This mechanism promotes sparse, decorrelated representations. The size of the input layer is determined either by the combined size of the EC populations or by the dimensionality of the MNIST images.

Parameters of both subnetworks are given in Table S1.

### The infrapyramidal blade network

The infrapyramidal blade network consists of the input population (EC) and two DG populations: excitatory and inhibitory (Fig. 1b), and is inspired by the model of Gozel and Gerstner (2021). Neurons are rate-based and the firing rate evolves according to:

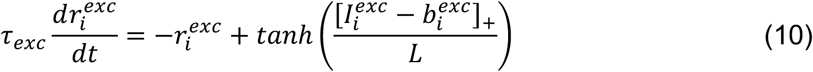

where τ*_exc_* is a time constant of the excitatory DG neurons, *I^exc^* is the input current, *L* is a smoothing parameter, b*^exc^* is the firing threshold, *r_i_* is the firing rate of excitatory neuron *i*.

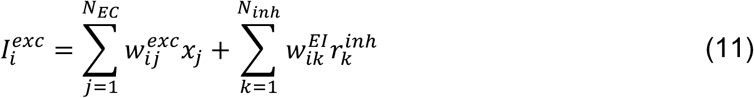

The inhibitory neurons evolve according to:

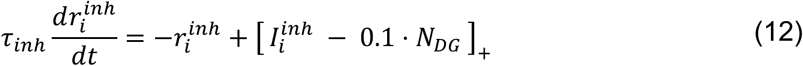

where τ*_inh_* is the inhibitory time constant, 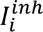 is the input current, 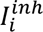 is the firing rate of inhibitory neuron *i*. A fixed offset equal to 0.1*N*_DG_ was applied to the inhibitory input current; with *N*_DG_ = 500, this offset was 50.

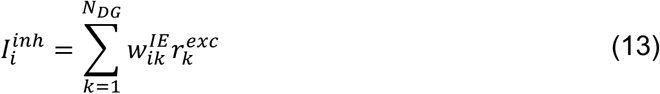

The network is trained using a heterosynaptic learning rule (Chistiakova et al., 2014; Zenke and Gerstner, 2017) as employed by Gozel and Gerstner (2021):

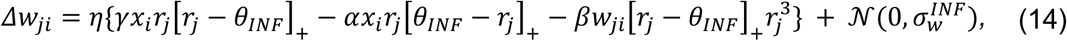

where *w*_j*i*_ is synaptic weight from neuron *i* to neuron *j*, *η* refers to the learning rate, *x_i_* is the activity of the input neuron, *r*_j_ the firing rate of the DG (postsynaptic) neuron, *θ_INF_* the firing threshold, *γ* = *γ*_0_ − *θ_INF_* indicates the strength of the LTP, 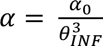 designates the strength of the LTD, *β* is the contribution of the heterosynaptic LTD, and 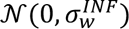 is the weight noise with the standard deviation 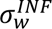. Heterosynaptic LTD refers here to the weight decrease of a synapse when the presynaptic neuron is not active but the postsynaptic neuron is active. Square brackets denote the positive-part operator, so only positive values contribute to this term.

The bias was updated once after each input-pattern presentation according to

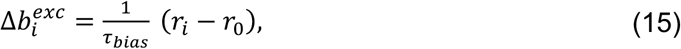

where *τ_bias_* is the bias-adaptation scale controlling the update magnitude per pattern presentation, *r_i_* is the excitatory DG firing rate, and *r*_0_ is the target firing rate. All network parameters for DG_SUP_ and DG_INF_ are listed in Table S1.

### GABA-switch

In the GABA-switch experiment, a subset of dentate gyrus (DG) neurons with weak input connectivity is replaced by newborn units whose feedforward weights are initialized to zero. At the onset of the early maturation phase, these neurons cannot initially be driven by entorhinal cortex. Instead, they receive indirect depolarizing input mediated by interneurons. For the plastic neurons, inhibitory-to-excitatory weights were set to their positive pre-switch magnitude, 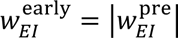, whereas reciprocal excitatory-to-inhibitory connections were set to zero, 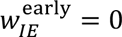, effectively converting GABAergic signaling into a depolarizing drive, corresponding to the early phase of the GABA-switch (Ge et al., 2006). Due to their increased excitability (*b^exc^* = 0) and lack of feedback projections, newborn neurons are co-activated with mature DG cells, leading to heterosynaptic growth of feedforward weights toward the average of currently represented inputs. In the subsequent late phase, the original inhibitory sign and magnitude were restored, 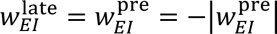, and reciprocal connections with interneurons were established (E-to-I connections were reinstated using a Bernoulli mask with probability 0.9; present connections had weight 1), introducing competition. The restoration of inhibitory connectivity corresponds to the progressive recruitment of feedback inhibition during maturation of abGCs (Temprana et al., 2015).

Together with increasing firing thresholds, now updated according to Eq. (15), this drives specialization of newborn neurons toward input patterns that are not yet well represented by the mature population. This two-phase mechanism implements a transition from cooperative integration to competitive pattern separation, enabling selective encoding of new inputs drawn from a related distribution.

### Training

For analyses using MNIST inputs, the two subnetworks were trained on 500 images, comprising 250 images from each of digit classes 3 and 4. The DG_SUP_ subnetwork was trained for one epoch using the fixed input order. The DG_INF_ subnetwork was trained for 20 epochs; the first epoch used the fixed input order, whereas all training patterns were randomly permuted during each of the remaining 19 epochs.

For analyses using the single-environment simulated EC dataset, the subnetworks were trained on five episodes in a square 2D environment. We sampled a 40 × 40 uniform grid and traversed adjacent grid points in a continuous serpentine sweep starting from the lower-left sampled location, yielding 1,600 spatially varying input patterns per episode and 8,000 patterns in total. The DG_SUP_ subnetwork was trained for one epoch with patterns ordered by episode and spatial position. The DG_INF_ subnetwork was trained for 10 epochs; the first epoch followed the original episode and position order, whereas all 8,000 training patterns were randomly permuted during each of the remaining nine epochs. Shuffling was used only for additional training passes and did not define a new physical trajectory.

For Figure 3c and 3d and Figure S5, the subnetworks are trained on 5 episodes in a square 2D environment and 5 episodes in a circular 2D environment. We use a 10×10 grid and retain only positions that exist in both environments. Because the circular environment has radius 1 m, this yields 88 shared positions out of the 100 grid positions. The DG_SUP_ subnetwork is again trained for 1 epoch with ordered inputs. The DG_INF_ subnetwork was trained for 50 epochs; the first epoch followed the original environment, episode, and position order, whereas the combined set of 880 patterns was randomly permuted during each of the remaining 49 epochs. The larger number of epochs partly compensated for the smaller number of input patterns relative to the single-environment dataset.

For Figures 6, S6, S7, and S8, the trained networks from the square-environment dataset are used as the basis for novel encoding. For MNIST, novel encoding used patterns from a novel digit class that was not included during initial training. For simulated EC inputs, we distinguish between novel same-environment and novel distinct-environment episodes. The novel same-environment episode was selected as a challenging held-out episode from the same environment distribution, defined as the episode with the largest object-position displacement relative to the training episodes, excluding the first five episodes used for training. For the novel distinct-environment episode, we used an episode from a circular environment.

## Analysis

### Positional tunings

Positional tunings were computed by dividing the 2 m × 2 m arena into a 40 × 40 grid. For each DG neuron, activity in each spatial bin was accumulated and divided by occupancy, with empty-bin counts set to 1 to avoid division by zero, yielding a raw firing-rate map. These maps were then smoothed with a Gaussian filter (*σ* = 1 bin) and normalized by their maximum value.

### Pattern similarity and separability

We quantify and visualize representational similarity using cosine similarity between neural activation patterns:

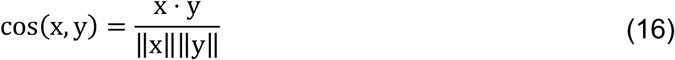

These similarities are organized into maps or matrices across positions or episodes and plotted as heatmaps. For Figure 4, pattern separability is defined as

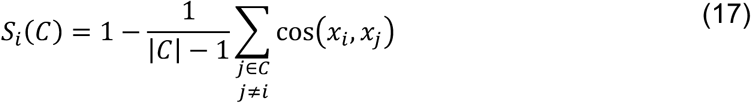

where *C* is the relevant comparison set.

### Place field size

Place field size was estimated from the smoothed positional tuning maps. For each neuron, a place field was considered valid only if the peak activity reached at least 0.2. The field threshold was then defined as the larger of 20% of that neuron’s peak activity and an absolute threshold of 0.2. Place field size was computed as the total area of all spatial bins whose activity exceeded this threshold. With a 2 m × 2 m arena tessellated into a 40 × 40 grid, each bin corresponded to (2/40)^2^ = 0.0025 m², so field size was obtained by multiplying the number of suprathreshold bins by the bin area.

### Weighted receptive fields

To summarize which input features contributed to a given DG response, we computed a weighted receptive field for each pattern as the activity-weighted average of the DG feedforward weights. Specifically, if *a* is the DG activity vector evoked by an input pattern and *W* is the matrix of feedforward receptive fields, the weighted receptive field was computed as

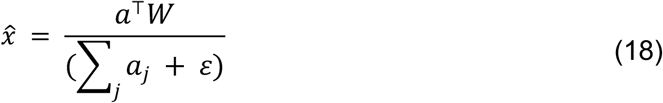

with *ε* = 10^-8^. This yields a pattern-specific average receptive field that emphasizes the neurons contributing most strongly to the response.

### Remapping on linearized trajectories

A linear trajectory for Figure 3 was selected according to two criteria: (1) it should include bins present in both the square and circular environments, enabling comparison across geometries; and (2) it should avoid bins that are spatially close to trajectory points traversed in the opposite direction. Figure S1b shows the selected trajectory together with the boundaries of the square and circular arenas. For each pairwise comparison, we first retained the union of neurons whose maximum activity along the trajectory exceeded 0.05 in at least one of the two conditions. In the reference condition, episode 1 for within-environment comparisons and environment A for between-environment comparisons, retained neurons exceeding 0.05 were sorted by the position of their maximum activity.

Neurons that exceeded the threshold only in the comparison condition retained their relative row positions within the filtered set. The resulting neuron ordering was applied unchanged to the comparison condition. Gaussian smoothing with *σ* = 1 bin was applied along the trajectory after sorting for visualization.

### Encoding and forgetting

For both the MNIST and neural experiments, we first fit linear ridge decoders from DG activity back to the input space separately for DG_SUP_ and DG_INF_, using only the patterns employed during initial training. Novel inputs consisted of either patterns from a novel MNIST digit class (here, “5”), a novel same-environment episode from the square-environment distribution, or a novel distinct-environment episode from the circular environment. These inputs were identified as poorly reconstructed by the decoder, i.e. inputs producing large prediction errors, with reconstruction similarity below the blade-specific threshold *th_enc_*. For DG_SUP_, novel encoding was restricted to a plastic subset of neurons (*N_imm_*), preselected from uncommitted neurons when available, or otherwise from neurons with the weakest feedforward weights. Before encoding, these neurons were reset to allow recruitment by the new inputs. The model was then trained only on the selected novel patterns, and the decoder was refit on the combined set of old and newly encoded patterns. For DG_INF_, novel encoding was likewise restricted to a weakly connected plastic subset, but learning followed a two-stage inhibitory-switch procedure. In the early phase, feedforward weights and biases of the selected plastic neurons were reset and inhibitory couplings were transiently reconfigured to facilitate recruitment by the novel patterns. In the late phase, inhibitory connectivity was restored to its normal sign and sparsity structure, and training continued for one epoch on the selected novel patterns alone. The early and late GABA-switch phases each comprised one epoch. After the late phase, the decoder was refit on the combined set of old and newly encoded patterns. Forgetting was applied only to DG_SUP_. Candidate old memories were selected either by explicit pattern identity for MNIST or, for simulated EC inputs, by selecting all patterns from one predefined trained episode. Within this candidate set, forgetting was triggered only for patterns whose DG_SUP_ and DG_INF_ decoder reconstructions both exceeded the preset decoder threshold 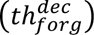, indicating that the memory was reliably represented in both streams. Among the active DG_SUP_ units for that pattern, neurons whose activity exceeded the DG_SUP_ activity threshold 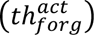 were deleted by zeroing their feedforward receptive fields and lateral inhibitory connections, and by resetting their commitment state, analogous to neuronal turnover. After forgetting, the DG_SUP_ decoder was refit on the complete original old-pattern set using post-forgetting DG_SUP_ activities. Exact parameter values are given in Table S2.

## Acknowledgements

VS is supported by the Max Planck School of Cognition. CFD’s research is supported by the Max Planck Society, the European Research Council, the Kavli Foundation, the Jebsen Foundation, Helse Midt Norge and The Research Council of Norway. AB acknowledges funding from the Max Planck Society. We thank all members of the neural computation group for fruitful discussions and feedback.

## Code availability

The code supporting the findings of this study will be made available after publication under https://github.com/Bicanski-NCG.

## Supplementary materials

### A computational model of the two dentate gyrus blades

**Table S1.** Model parameters.

| Parameter | MNIST - DG<sub>SUP</sub> | MNIST - DG<sub>INF</sub> | Simulated neural inputs - DG<sub>SUP</sub> | Simulated neural inputs - DG<sub>INF</sub> |
| --- | --- | --- | --- | --- |
| $N_{EC}$ | 144 | 144 | 1003 | 1003 |
| $N_{DG}$ | 500 | 500 | 500 | 500 |
| $\eta$ | 1 | 0.01 | 1 | 0.01 |
| $N_{DG}^{inh}$ | - | 100 | - | 100 |
| $w_{IE}$ | - | 1 | - | 1 |
| $w_{EI}$ | - | $-1/(p_{EI} * N_I)$ | - | $-1/(p_{EI} * N_I)$ |
| $p_{IE}$ | - | 0.9 | - | 0.9 |
| $p_{EI}$ | - | 0.9 | - | 0.9 |
| $\tau_{exc}$ | - | 20 ms | - | 20 ms |
| $\tau_{inh}$ | - | 2 ms | - | 2 ms |
| $\tau_{bias}$ | - | 100 | - | 100 |
| $r_0$ | - | 0.1 | - | 0.1 |
| $L$ | - | 0.5 | - | 0.5 |
| $\alpha_0$ | - | 0.05 | - | 0.05 |
| $\beta$ | - | 1.5 | - | 1.5 |
| $\gamma_0$ | - | 10 | - | 10 |
| $\theta_{INF}$ | - | 0.15 | - | 0.15 |
| $\Delta t$ | - | 0.1 ms | - | 0.1 ms |
| $\theta_{SUP}$ | 0.95 | - | 0.88 | - |
| $\eta_{reuse}$ | 0.3 | - | 0.3 | - |
| $\eta_{inh}$ | 0.1 | - | 0.1 | - |
| $\lambda_{inh}$ | 0.001 | - | 0.001 | - |
| $L_{max}$ | 2.0 | - | 2.0 | - |
| $\sigma_{SUP}$ | 0.03 | - | 0.05 | - |
| $\sigma_w^{INF}$ | - | 0.005 | - | 0.01 |
| k | 10 | - | 10 | - |
| *Dataset size* | 500 | 500 | 8000 | 8000 |
| *Training epochs* | 1 | 20 | 1 | 10 |

Simulated-neural values refer to the single-environment models. For the two-environment analyses in Figures 3c–d and Figure S5, the combined dataset contained 880 patterns, DG_SUP_ used *θ*_SUP_ = 0.65 and one training epoch, and DG_INF_ used 50 training epochs.

**Table S2.** Parameters for encoding and forgetting experiments in Figures 5 and 6.

| Parameter | MNIST - $DG_{SUP}$ | MNIST - $DG_{INF}$ | Simulated neural inputs - $DG_{SUP}$ | Simulated neural inputs - $DG_{INF}$ |
| --- | --- | --- | --- | --- |
| $th_{enc}$ | 0.85 | 0.85 | 0.85 | 0.85 |
| $th_{forg}^{dec}$ | 0.85 | 0.85 | 0.85 | 0.85 |
| $th_{forg}^{act}$ | 0.85 | - | 0.85 | - |
| $\lambda_{dec}$ | 0.01 | 0.01 | 0.01 | 0.01 |
| $N_{imm}$ | 50 | 50 | 100 | 100 |
| *Early GABA-switch epochs* | - | 1 | - | 1 |
| *Late GABA-switch epochs* | - | 1 | - | 1 |

**Table S3.** Parameter descriptions.

| Parameter | Descriptions |
| --- | --- |
| $N_{EC}$ | input size |
| $N_{DG}$ | size of the DG cell population |
| $\eta$ | learning rate |
| $N_{DG}^{inh}$ | size of the DG inhibitory cell population |
| $w_{IE}$ | weights from excitatory DG cells to inhibitory DG cells |
| $w_{EI}$ | weights from inhibitory DG cells to excitatory DG cells |
| $p_{IE}$ | fraction of excitatory DG-to-inhibitory interneuron connections |
| $p_{EI}$ | fraction of inhibitory interneuron-to-excitatory DG connections |
| $\tau_{exc}$ | membrane time constant for DG activity dynamics |
| $\tau_{inh}$ | time constant for inhibitory dynamics |
| $\tau_{bias}$ | bias-adaptation scale controlling the update magnitude per pattern presentation |
| $r_0$ | target firing rate for all $DG_{INF}$ neurons |
| $L$ | slope or gain parameter of the $DG_{INF}$ output nonlinearity |
| $\alpha_0$ | activity-dependent depression coefficient in the heterosynaptic learning rule |
| $\beta$ | strength of heterosynaptic competition or weight decay in the $DG_{INF}$ rule |
| $\gamma_0$ | how strongly weights increase when postsynaptic activity exceeds threshold |
| $\theta_{INF}$ | postsynaptic firing threshold used in the $DG_{INF}$ heterosynaptic plasticity rule |
| $\Delta t$ | integration step size for the rate-based dynamics |
| $\theta_{SUP}$ | similarity threshold used to determine whether an input matches an existing exemplar and whether winner neurons are eligible for the exemplar reuse update |
| $\eta_{reuse}$ | step size for the exemplar integration/reuse update |
| $\eta_{inh}$ | learning rate for lateral inhibition between exemplar neurons |
| $\lambda_{inh}$ | decay applied to previously learned lateral inhibition weights |
| $L_{max}$ | upper bound on learned lateral inhibition strength |
| $\sigma_{SUP}$ | width parameter controlling the soft similarity-to-rate activity readout in $DG_{SUP}$ |
| $\sigma_w^{INF}$ | standard deviation of $DG_{INF}$ weight noise |
| $k$ | number of winner neurons selected during $DG_{SUP}$ k-WTA competition |
| *Dataset size* | number of patterns in the input dataset |
| *Training epochs* | number of complete passes through the training dataset |
| $th_{enc}$ | blade-specific decoder reconstruction cosine-similarity threshold below which an input is selected for novel encoding |
| $th_{forg}^{dec}$ | minimum reconstruction cosine similarity that must be reached by both the $DG_{SUP}$ and $DG_{INF}$ decoders for a pattern to be eligible for forgetting |
| $th_{forg}^{act}$ | minimum $DG_{SUP}$ activity required for a neuron to be selected for deletion during forgetting |
| $\lambda_{dec}$ | ridge regularization coefficient used when fitting the linear decoders |
| $N_{imm}$ | number of plastic neurons assigned to the neurogenesis-inspired turnover subset during novel encoding |

**Figure S1.**
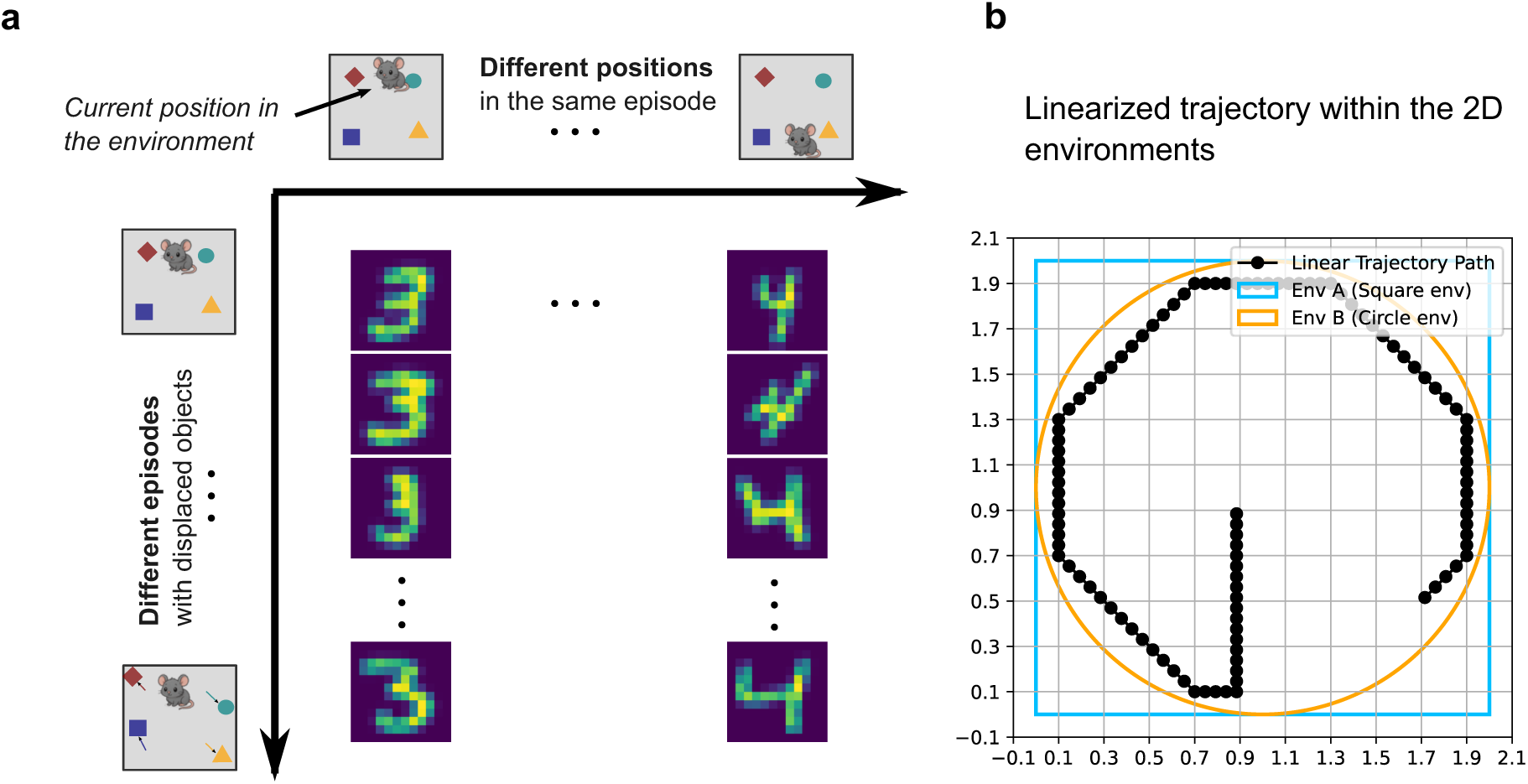
Analogy between MNIST and simulated neural inputs. **a:** Conceptual correspondence between MNIST inputs and simulated spatial episodes. Different handwritten instances of the same digit class are treated as analogous to repeated visits to the same environment, where the agent occupies corresponding positions but object locations and random spatial cell activity vary across episodes. In both cases, individual inputs correspond to episodes, whereas averages across related inputs correspond to integrated prototypes. **b:** Illustration of the linearized trajectory used to sample activity from the 2D simulated environment. Each trajectory position corresponds to the agent’s current location and is used to compare activity patterns across positions and episodes.

**Figure S2.**
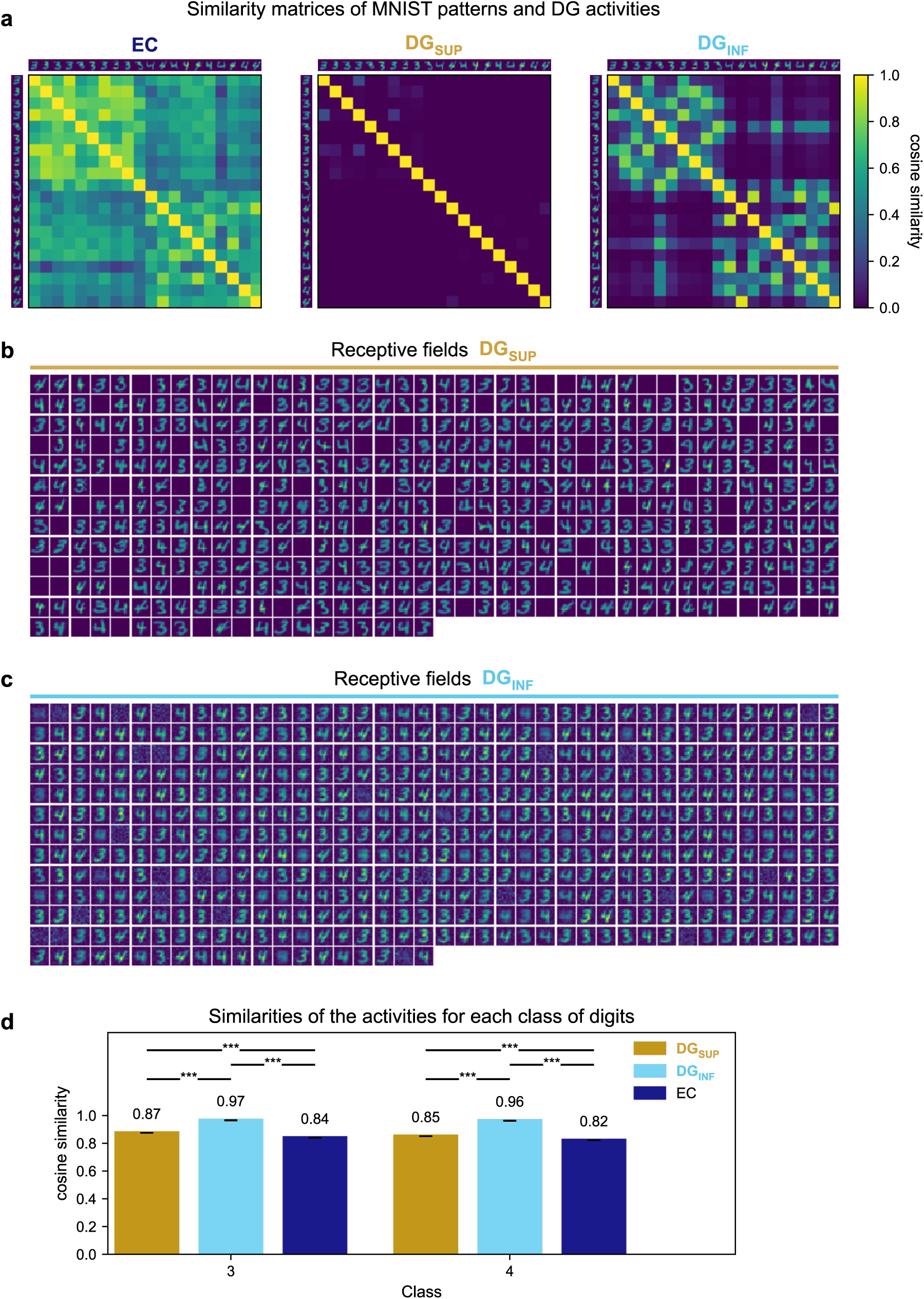
Similarity structure and receptive-field organization in MNIST representations. **a:** Pairwise cosine similarity matrices for 20 selected MNIST digits from classes 3 and 4 in EC, DG_SUP_, and DG_INF_. **b–c:** Normalized receptive fields of DG_SUP_ **(b)** and DG_INF_ **(c)** neurons. **d:** Cosine similarity between activity-weighted receptive-field reconstructions and the corresponding class-mean image. DG_INF_ representations were more similar to the class mean than DG_SUP_ representations for both digit classes, consistent with prototype-like integration. For class 3: DG_SUP_, 0.87, SD < 0.001; DG_INF_, 0.97, SD = 0.001; EC, 0.84. For class 4: DG_SUP_, 0.85, SD < 0.001; DG_INF_, 0.96, SD = 0.001; EC, 0.82. Values are reported as means with SDs across 50 independently seeded training runs per blade; EC is a fixed input-derived baseline. DG_SUP_–DG_INF_ comparisons used two-sided Welch independent-samples t-tests, whereas comparisons against the fixed EC baseline used two-sided one-sample Wilcoxon signed-rank tests. Holm correction was applied across the six prespecified classwise comparisons. ***Adjusted p < 0.001.

**Figure S3.**
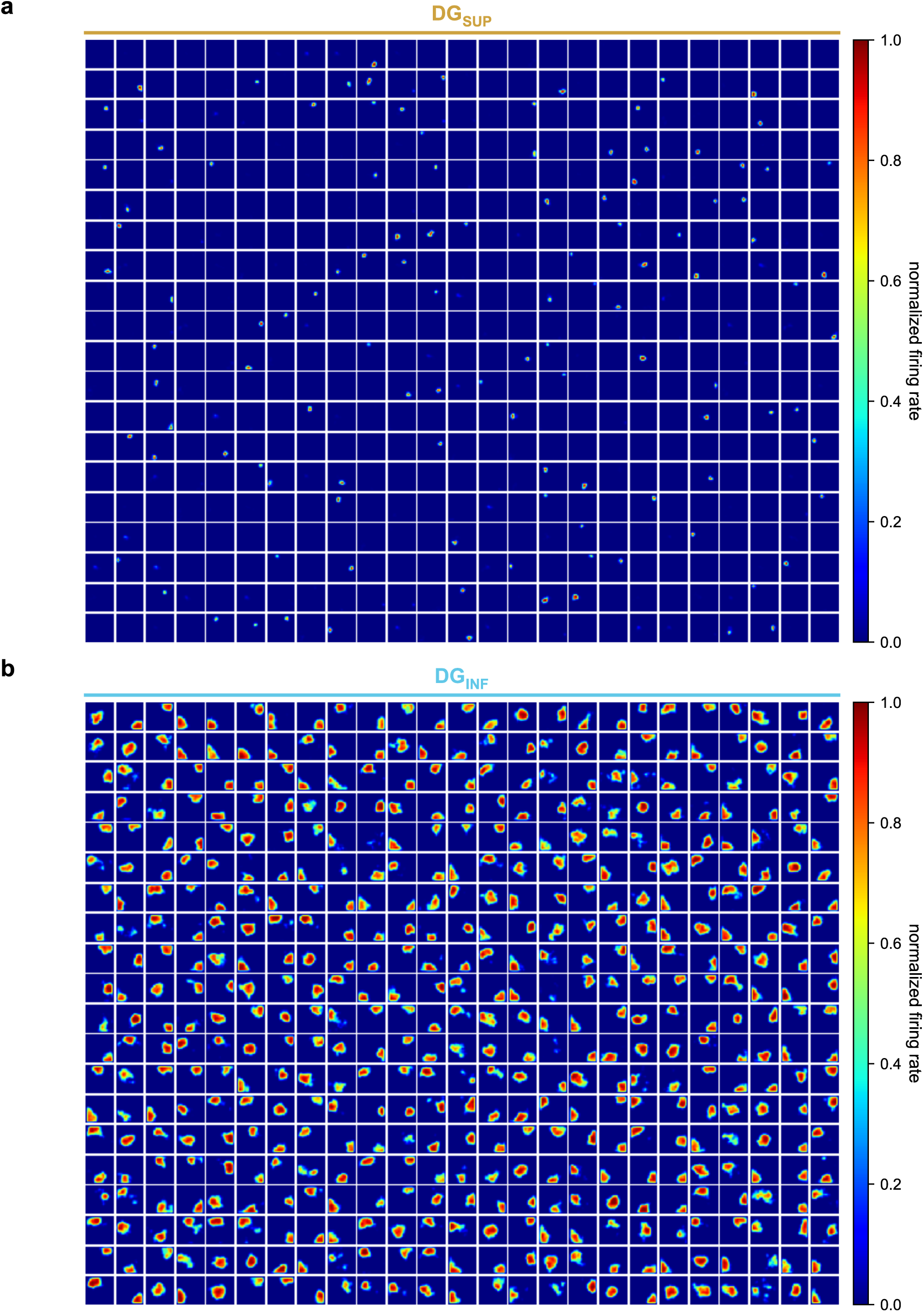
Positional tuning maps of DG_SUP_ and DG_INF_ neurons trained on simulated EC inputs. **a:** Positional tuning maps of all 500 DG_SUP_ neurons in a representative episode after training on five episodes. DG_SUP_ neurons show sparse and spatially localized firing fields. **b:** Positional tuning maps of all 500 DG_INF_ neurons after training on the same inputs. Compared with DG_SUP_, DG_INF_ neurons show broader positional tuning, consistent with integration across similar episodes. All maps are shown on a normalized firing-rate scale.

**Figure S4.**
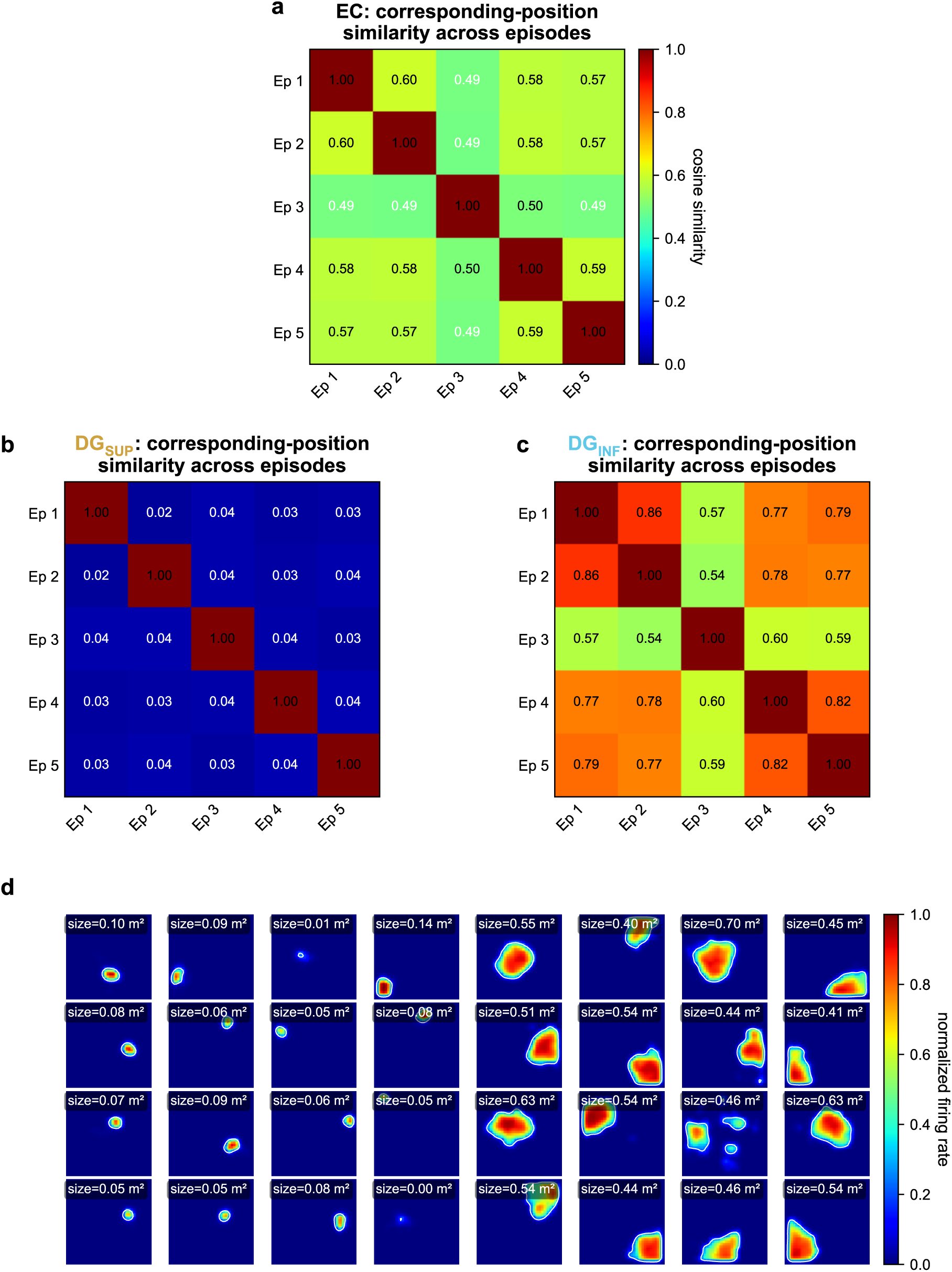
Similarity structure and place field detection for simulated EC inputs. **a-c:** Similarity matrices for EC **(a)**, DG_SUP_ **(b)**, and DG_INF_ **(c)** activity patterns across episodes. Matrix values indicate mean pairwise cosine similarity, with higher values reflecting more stable representations across episodes. DG_SUP_ representations show low similarity across episodes, whereas DG_INF_ representations preserve higher similarity, consistent with integration across similar experiences. **d:** Example positional tuning maps from DG_SUP_ and DG_INF_ used for place field size detection. Maps are shown on a normalized firing-rate scale, and white contours mark the detected place field area.

**Figure S5.**
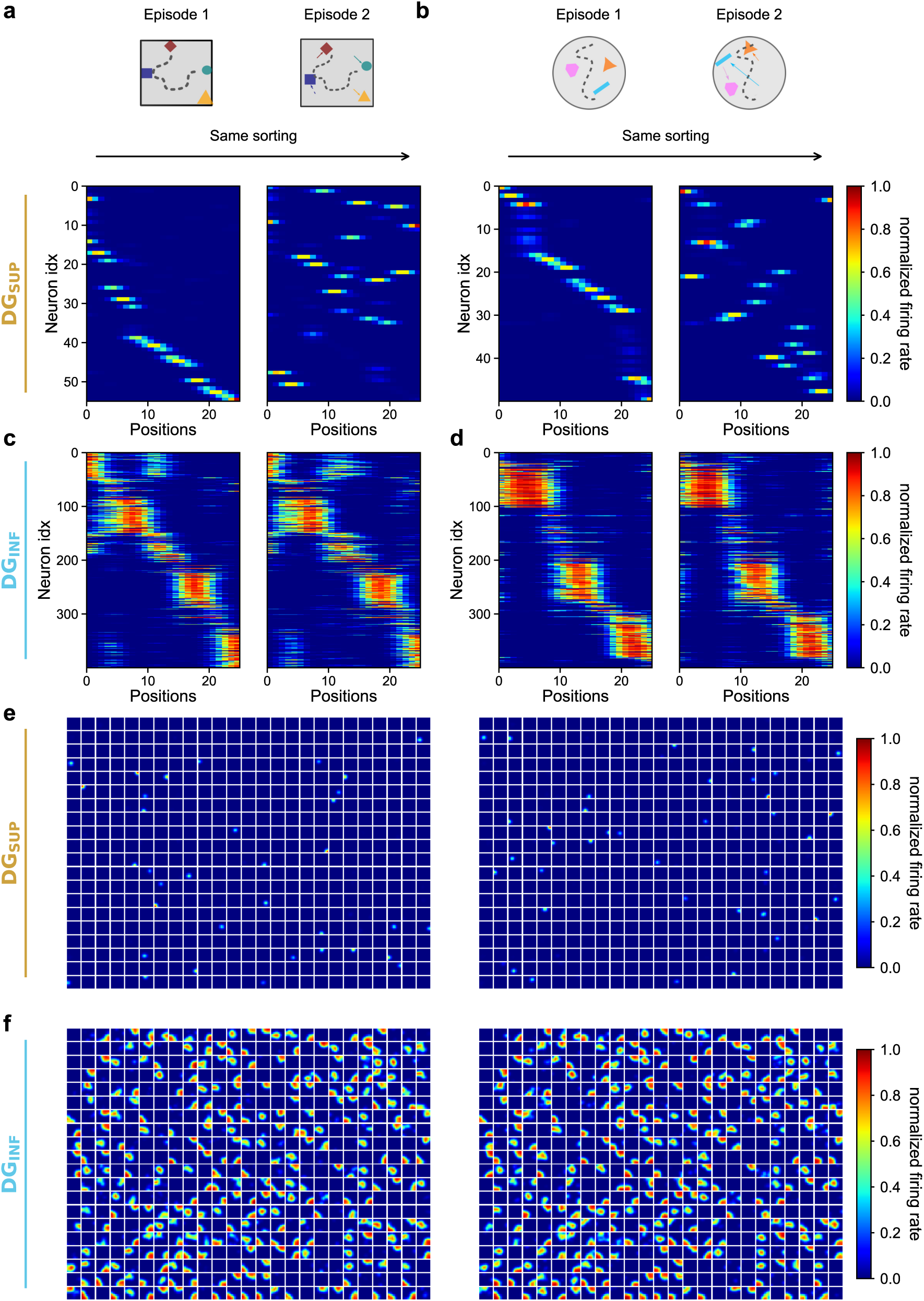
Within-environment control for blade-specific place field stability and remapping along linearized trajectories. **a-b:** DG_SUP_ activity along linearized trajectories in two episodes from the same environment. Neurons are sorted by their peak firing position in episode 1, and the same ordering is applied to episode 2. DG_SUP_ neurons show pronounced episode-specific remapping within both the square environment **(a)** and the circular environment **(b)**. **c–d:** DG_INF_ activity shown with the same procedure. In contrast to DG_SUP_, DG_INF_ neurons preserve a largely stable diagonal structure across episodes within both the square environment **(c)** and the circular environment **(d)**, consistent with integration across similar experiences. **e–f:** Two-dimensional positional tuning maps for the same within-environment episode comparison. DG_SUP_ tuning maps change substantially between episode 1 and episode 2 **(e)**, whereas DG_INF_ tuning maps remain largely stable across episodes **(f)**, matching the remapping and stability observed along the linearized trajectories. All panels use the same subsampled 10 × 10 spatial grid as in Fig. 3c–d. Only neurons whose activity exceeds 0.05 in at least one of the two compared episodes are plotted, thereby retaining cells active in either episode while excluding cells silent in both.

**Figure S6.**
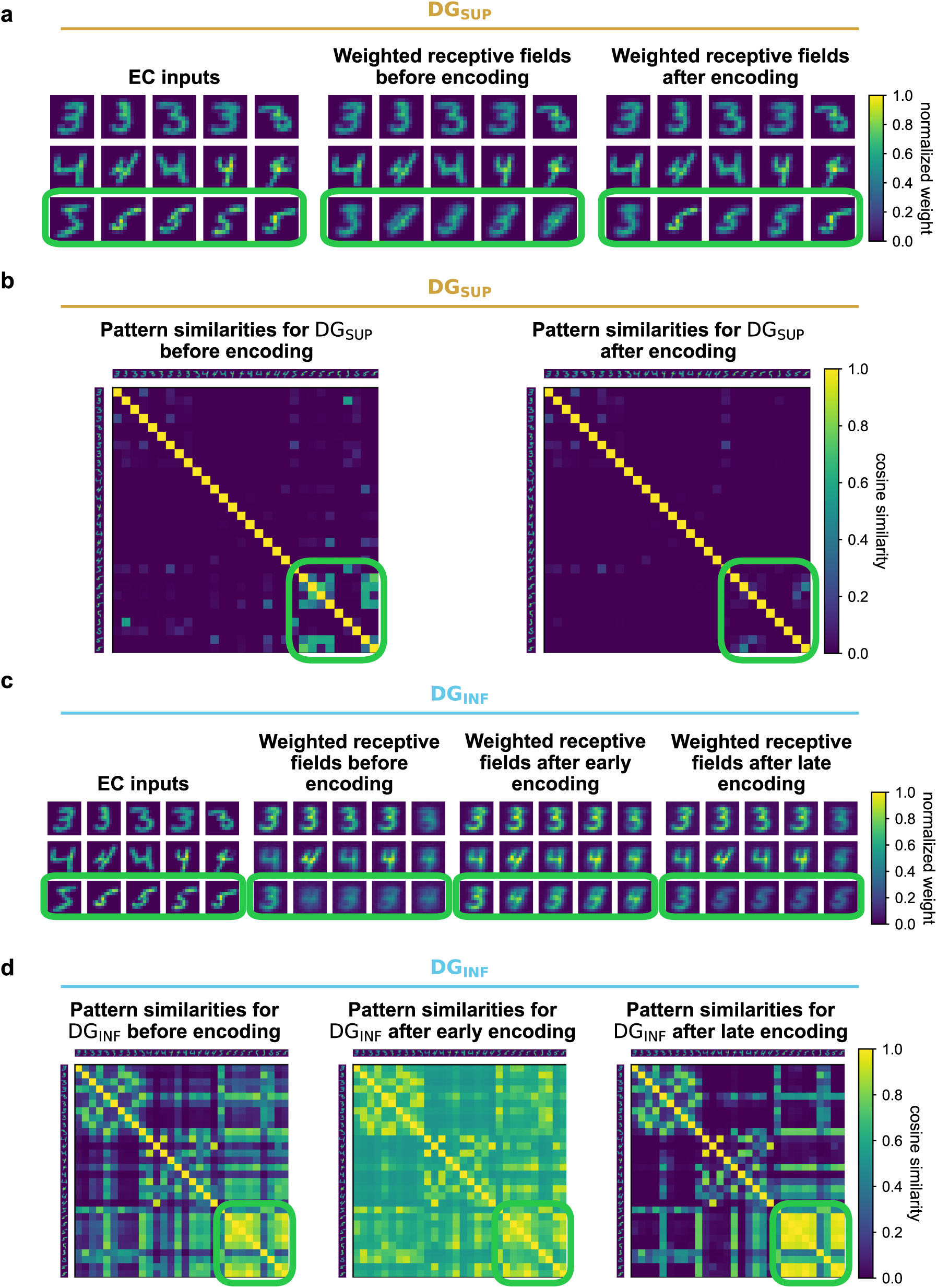
Encoding dynamics of novel MNIST class in DG_SUP_ and DG_INF_. **a:** Novel-class encoding in DG_SUP_. A subset of EC input patterns is shown together with the corresponding activity-weighted DG_SUP_ receptive fields before and after encoding. Newly recruited neurons acquire receptive fields for novel digit-class exemplars after one-shot learning (highlighted in green). **b:** Pairwise cosine similarity between DG_SUP_ representations before and after encoding. After encoding, novel-class patterns become more separated from previously stored representations, consistent with exemplar-like encoding. **c:** Novel-class encoding in DG_INF_. The same input patterns are shown together with activity-weighted DG_INF_ receptive fields before encoding, after the early GABA-switch phase, and after the late GABA-switch phase. **d:** Pairwise cosine similarity between DG_INF_ representations across the same encoding stages. Similarity increases during the early phase, reflecting integration of novel-class patterns, and is partially refined after the late phase.

**Figure S7.**
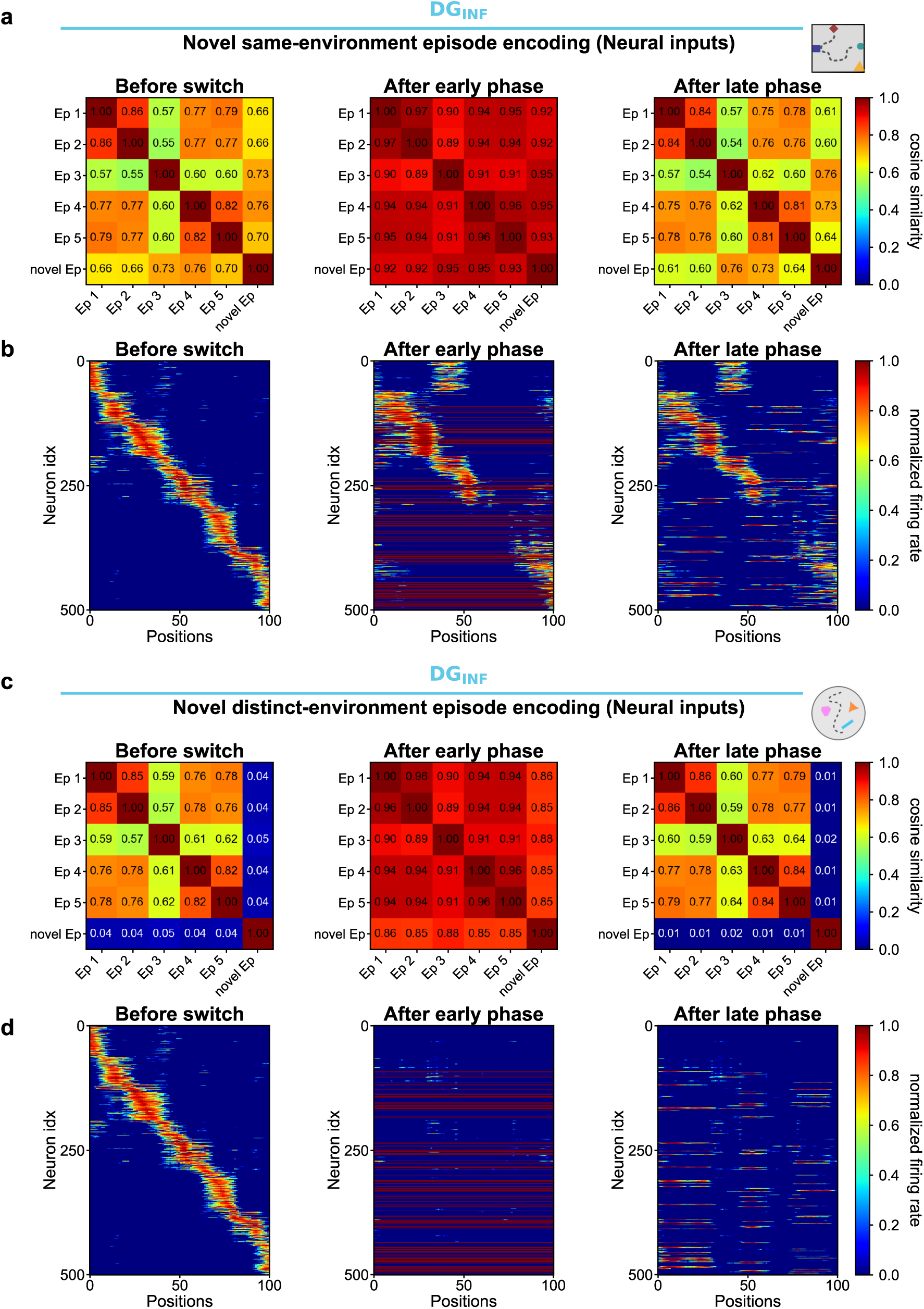
Encoding dynamics for simulated EC inputs in DG_INF_. **a:** Episode-similarity matrices before encoding of a novel same-environment episode, after the early GABA-switch phase, and after the late GABA-switch phase. The novel same-environment episode has the same environment geometry and object identities as the training episodes, but a different object constellation. **b:** DG_INF_ activity along a linearized trajectory for the same novel same-environment episode comparison. Neurons are sorted by peak firing position in a previously encoded episode, and the same ordering is applied after early and late GABA-switch encoding. The early phase increases integration across episodes, whereas the late phase refines positional tuning while preserving similarity between related episodes. **c:** Episode-similarity matrices for encoding of a novel distinct-environment episode with different environment geometry, shown before encoding and after the early and late GABA-switch phases. **d:** DG_INF_ activity along a linearized trajectory for the novel distinct-environment comparison. After the early phase, activity is broadly integrated, whereas after the late phase DG_INF_ forms a distinct representation for the novel environment. All firing-rate maps are shown on a normalized color scale.

**Figure S8.**
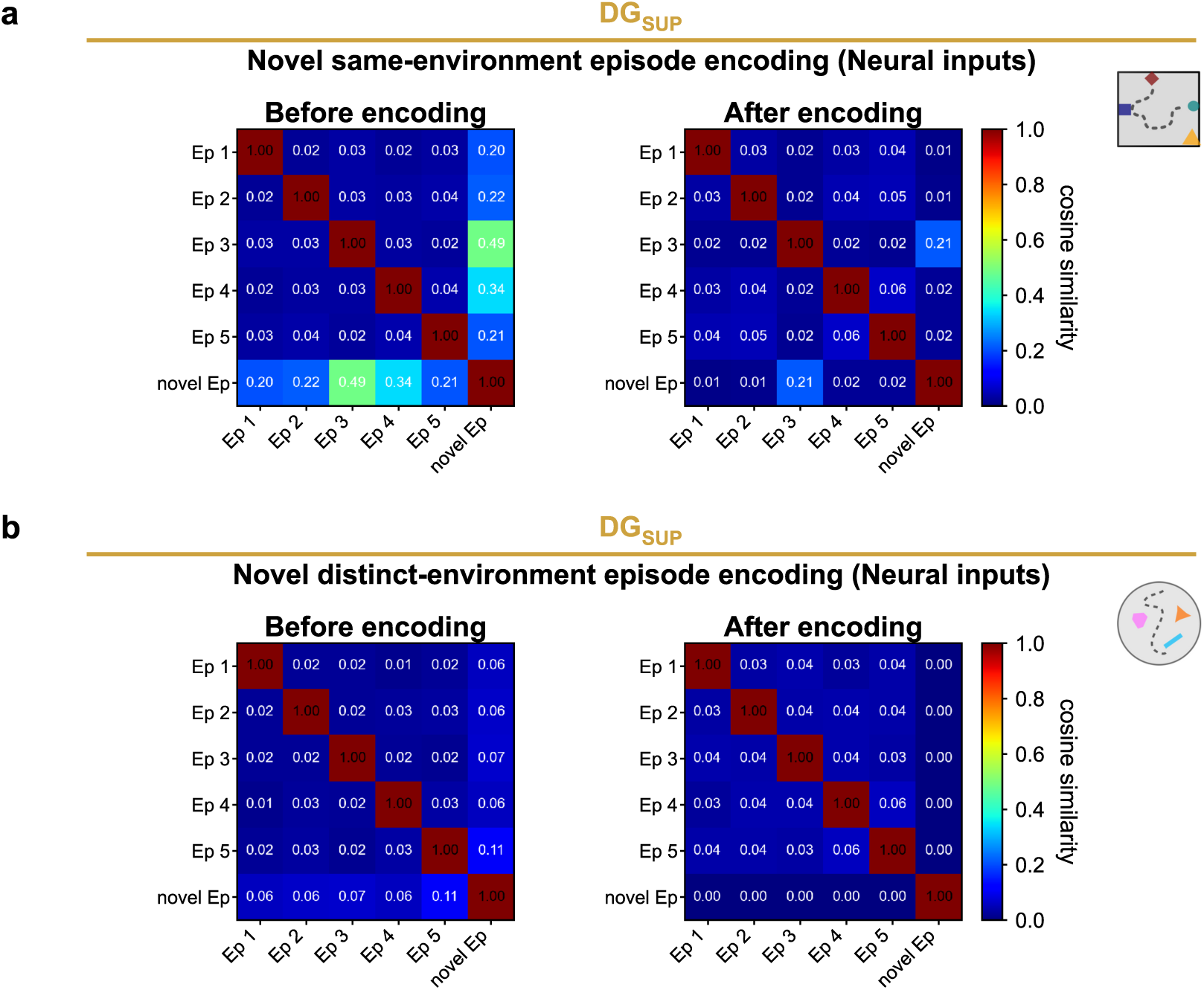
Encoding dynamics for simulated EC inputs in DG_SUP_. **a:** Episode-similarity matrices for DG_SUP_ activity before and after encoding of a novel same-environment episode. The novel same-environment episode has the same environment geometry and object identities as the training episodes, but a different object constellation. After encoding, DG_SUP_ forms a separated representation for this episode, consistent with exemplar-like episodic encoding. **b:** Episode-similarity matrices for DG_SUP_ activity before and after encoding of a novel distinct-environment episode with different environment geometry. As for the novel same-environment episode, encoding produces an orthogonalized representation of the new episode, indicating strong pattern separation in DG_SUP_.

